# tugMedi: simulator of cancer-cell evolution for personalized medicine based on the genomic data of patients

**DOI:** 10.1101/2025.06.27.661855

**Authors:** Iurii Nagornov, Eisaku Furukawa, Momoko Nagai, Shigehiro Yagishita, Tatsuhiro Shibata, Mamoru Kato

## Abstract

Cancer comprehensive genomic profiling tests are increasingly used, but drug response rates remain limited. Simulations forecasting cancer progression could aid targeted therapies; however, existing simulations focus mainly on basic biology. We present *tugMedi*, a cancer-cell evolution simulator designed for cancer genome medicine. By integrating patient-specific genomic and imaging data, tugMedi reconstructs each tumor’s genomic features and growth behavior, enabling real-time predictions of clonal dynamics under virtual drug regimens. tugMedi explicitly models copy number alterations and SNVs on parental chromosomes in recessive and dominant modes, capturing loss-of-heterozygosity and yielding precise variant allele frequencies and tumor contents. It handles mutations in cancer-related genes with real exon-intron structures by a fast algorithm. For TCGA samples, it provided ensemble predictions of clonal dynamics, allowing extraction of drug response, tumor-size shrinkage rates, and time to recurrence under specific drug conditions. tugMedi represents a first step toward simulation-driven genome medicine based on a patient-derived virtual tumor.

## INTRODUCTION

Cancer genome medicine is being incorporated into clinical practice. Typically, in cancer comprehensive genomic profiling (CGP), DNA variants in >100–500 genes are detected in a single test by a medical device based on next-generation sequencing (NGS). Detected variants are targeted by molecular targeted drugs to inhibit the aberrant activities of the protein products or unlock suppressed immune activities. For example, DNA extracted from hospital-use, formalin-fixed paraffin-embedded samples derived from the cancer tissue of patients are clinically examined by NGS to detect SNVs/indels and copy number alterations (CNAs) in all exons of 114 genes and fusions of 12 genes in a single CGP test^1,2^. Detected variants and genomic data are systematically collected nationwide to suggest targeted drugs suitable against the variants and can be used to improve future targeted therapies^3^. However, current targeted therapies are only effective in a small fraction of patients.

It is of great benefit if a computer simulation predicts the time course of cancer states under targeted therapies determined by genomic medicine. Many algorithms and simulators for the evolution of cancer cells have been developed in cancer genomics based on NGS data^4–13^. However, such simulators have focused on basic biology, such as recapitulating intra-tumor heterogeneity (ITH). Few studies have addressed therapeutic intervention in the context of cancer genome medicine, beyond understanding ITH. Although we previously addressed it in a cancer-genomic evolution simulator in part^14^, existing simulators are insufficient for practical application.

To begin with, simulators need to be more realistic to simulate therapeutic intervention. For example, most simulators either omit CNAs or address them in simplified forms^4,15^, resulting in inaccurate representation of variant allele frequencies (VAFs) affected by CNAs. Few simulators consider recessive and dominant genetic modes, leaving loss-of-heterogeneity (LOH) unrepresented. Most simulation models also fail to precisely simulate multi-stage tumorigenesis due to the rarity of multiple occurrences based on the low mutation rate times short gene sizes, causing long waiting times. Additionally, simulators lack a real-time framework for therapeutic interventions. Although some aspects may be partially addressed, integrating these realistic components into a single simulator is essential.

Herein, we developed the *tugMedi* (*tu*mor-*g*enome *Medi*cal) simulator. tugMedi uses the genomic data and radiological imaging data of tumors obtained from individual patients to predict cancer evolutionary dynamics under different settings of targeted therapies.

## RESULTS

### Overview of tugMedi

The overview of tugMedi is shown in **Figure 1**; see **Methods** for the details. tugMedi simulates the dynamics of the clones of cells in a Markov-chain process, where clones refer to groups of cells with identical mutations. According to input parameters such as the cell division and mutation rates, the states of clones stochastically change with time and new clones may be generated. Passenger and driver mutations randomly occur in clones at random times generated from user-specified distributions, such as an exponential and uniform distributions, respectively (see EF algorithm in **Methods**; **Supplementary Figure 1**). Users can restrict driver mutations to occur only in clones with a driver mutation on a particular gene. For example, a *KRAS* mutation can be restricted to occur only in clones with a driver mutation in *APC*.

**Figure 1.**
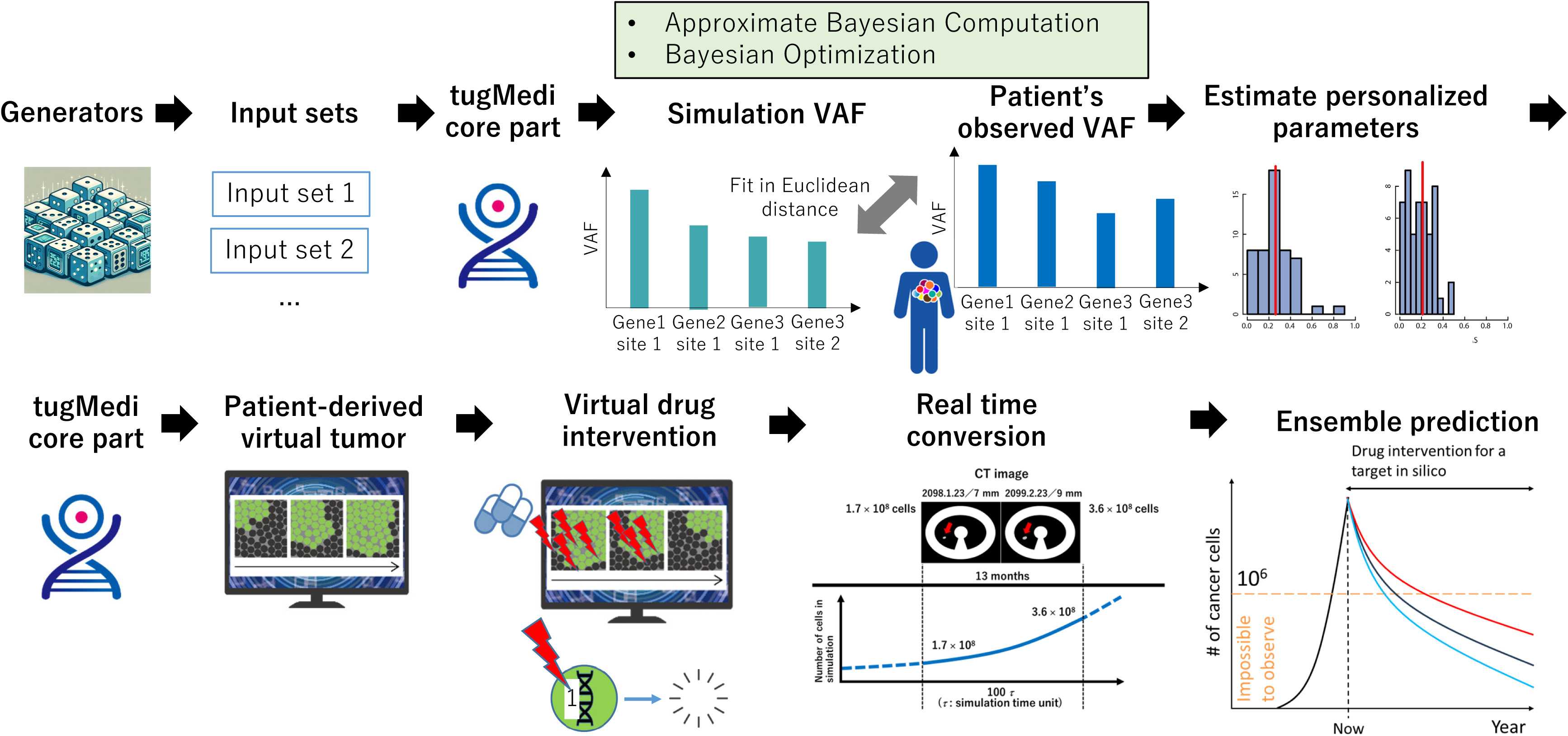
Simulation procedure until ensemble prediction. Generators generate random values for various parameters and multiple input sets are prepared. tugMedi executes simulations, and ABC and BO are performed to estimate the values of patient-specific simulation parameters. Based on estimated parameter values, tugMedi executes simulations to computationally grow a single cell to, say, 10^9^ cells. In the patient-derive virtual tumor, virtual drug intervention is performed to kill cells with aberrations in a targeted gene. Based on the change in tumor size according to patient’s radiological images, the simulation time unit is converted into the real time unit with a patient-specific conversion rate. Ensemble prediction is derived from multiple simulation replications. In the associated graph, the red, black, and blue curves represent the upper (e.g., 75^th^ percentile), average (median), and lower (25^th^ percentile) prediction values, respectively.

Point mutations, abbreviated as *poms*, correspond to SNVs/indels. They are distinguished from CNAs such as deletions (also referred to as loss) and duplications (amplifications or gains) (**Supplementary Figure 2**). Poms are treated as single points along chromosomes and occur only in exons, while CNAs are considered segments and occur in both exons and introns. The location information of the exons and introns of genes is obtained from the NCBI Consensus CDS database of the human reference genome. One breakpoint of a CNA randomly occurs anywhere in exons and introns (**Supplementary Figure 2**). The other breakpoint, or equivalently the CNA length is determined from a random number generated from an exponential distribution with the mean length derived from observed CNA lengths. CNA segments flanked by the breakpoints are removed from or duplicated in the chromosomes of clones for deletions or duplications, respectively (**Supplementary Figure 3**).

We refer to cells that acquire driver mutations as tumor/cancer cells. Driver mutations increase the cell division rate according to the weight parameter assigned to each driver gene, such as 0.2, 0.3, and 0.5 to *APC*, *KRAS*, and *TP53*, respectively. Users need to define specific driver genes (termed “regions of interest” in **Methods**). Weight parameters have to be estimated from observation data. Without driver genes, cells are usually set in the equilibrium state between cell division and cell death, with the cell division rate at zero on average^13^. The clones of cells with aberrant genes, that is, driver genes with driver mutations, have a net positive cell-division rate, and clones with more aberrant genes reproduce more stochastically (**Figure 2**).

**Figure 2.**
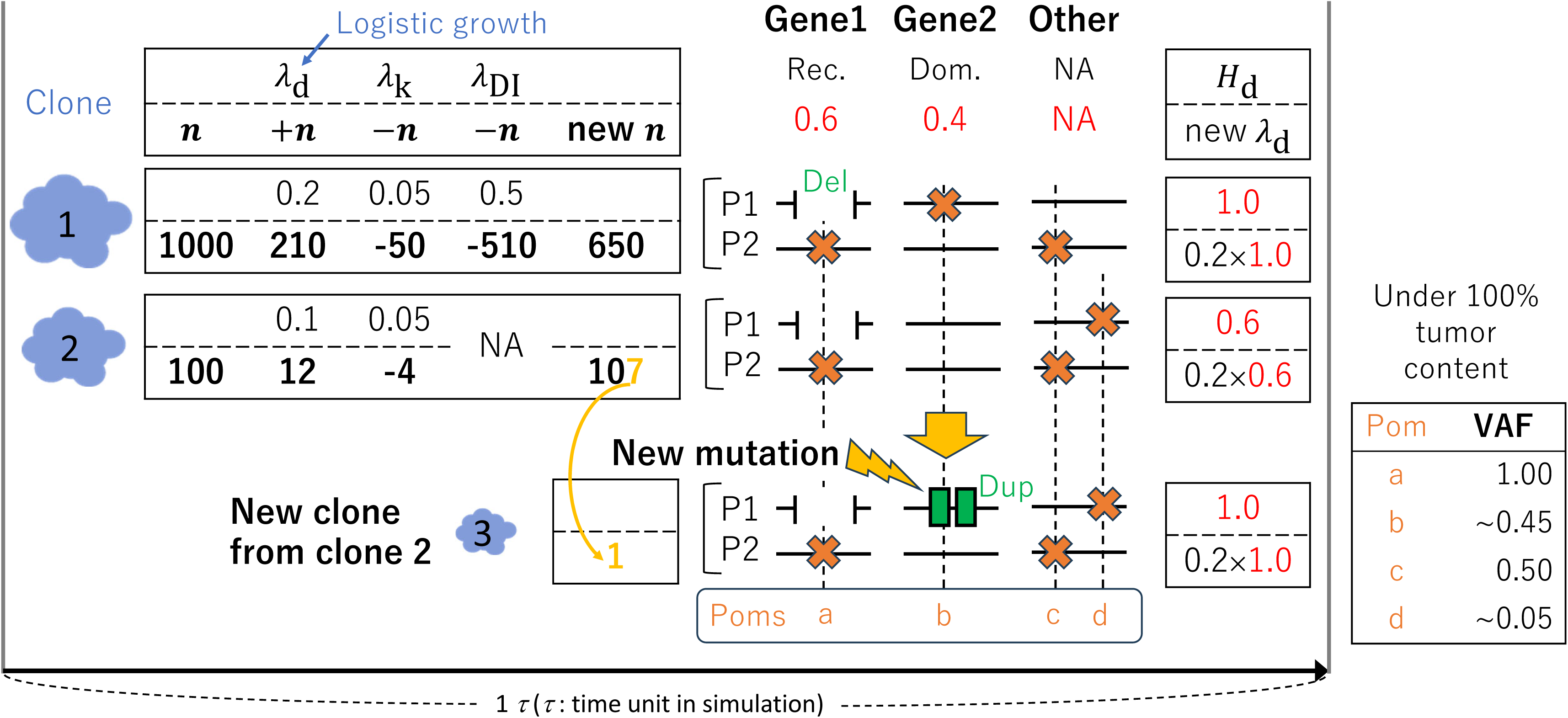
The central part of tugMedi. *n*, +*n*, —*n*, and new *n* are the number of cells, the increment, the decrement, and the number of cells for the next simulation timestep, respectively. The values of +*n* and —*n* are randomly determined from Poisson distributions with the means of *λ*_d_, *λ*_k_, and *λ*_DI_, for the trials of cell division, constant cell death, and drug intervention, respectively. *λ*_d_ is modelled to reflect logistic growth. Drug intervention targets only clones with aberration in gene2. Weight parameters are exemplified as 0.6 and 0.4 for gene1 and gene2, respectively, defined as driver genes. “Other” is defined as a region of non-driver genes, in which only passenger mutations occur, without a weight parameter. P1 and P2 represent parental chromosomes. A deletion and duplication occur in gene1 for clone1 and gene2 for clone3, respectively. The cross in orange represents point mutations, abbreviated as poms. Here, gene1 and gene2 are defined to follow the recessive (Rec.) and dominant (Dom.) models. A new clone, clone3 is generated from clone2 with a new mutation (duplication) at a timestep randomly specified from a prior distribution. *H*_d_ is the increment of the cell division rate calculated from the sum of the weight parameters of the aberrant driver genes, and converted into the Poisson parameter of new *λ*_d_ for the next timestep with a specified coefficient (here, 0.2). VAFs are calculated from the copy numbers of alleles in parental chromosomes, the fraction of subpopulations, and the normal-cell contamination. In this example, the normal-cell contamination (1 − tumor content) is assumed as 0.

In the simulator, parental chromosomes and the recessive and dominant models are implemented (**Supplementary Figure 4**). In recessive genes, driver mutations in *both* parental chromosomes result in an increased cell-division rate. Meanwhile, driver mutations in dominant genes in *either* parental chromosome yield the effect. Thus, LOH is expressed when one parental chromosome has a pom and the other has a deletion over the pom. VAFs from simulations are calculated using the copy numbers of alleles in parental chromosomes, the fraction of subpopulations, and the normal-cell contamination (**Supplementary Figure 5**), the last of which is ultimately estimated from observation data.

We used approximate Bayesian computation (ABC) and Bayesian optimization (BO) to estimate parameters (**Figure 1**). ABC estimates the posterior distributions by selecting parameter values for which simulated VAFs are close to observed VAFs. BO provides the optimal point estimate from a joint posterior distribution implied by parameter values selected in ABC. We implemented “generators” to generate random values for simulation parameters that users arbitrarily select, not limited to the cell-division weight parameters (**Supplementary Figure 6**). tugMedi takes in parameter values generated from generators. We implemented virtual drug intervention, where a virtual drug stochastically kills cells with user-specified aberrant genes at a user-specified killing rate per time, corresponding to a drug dose. This rate is mapped to a real dose via radiological imaging (**Supplementary Figure 7**). Values calculated in multiple simulation replications are finally summarized as percentiles at each timepoint and the percentiles are provided as ensemble predictions.

The simulation time unit is converted to a real time unit, in years and months, based on the time-change of the tumor size calculated from radiological image data (**Supplementary Figure 7**). Intra- and inter-tumor heterogeneity are reflected in the simulator from differences in simulation parameters estimated from patient-specific observed VAFs, and in the time-unit conversion rate derived from patient-specific imaging data.

### Simulation examples using synthetic data

In this section, we present simulations performed using synthetic data, especially focusing on the accuracy of estimation and prediction. The data were generated using pre-defined parameter values, referred to as the “true” values. For details on the data generation, see **Methods**.

#### 1) Synthetic data 1

This synthetic dataset was generated based on a lung cancer patient. In this dataset, we defined a short deletion in *TP53* and SNV in *BRAF* as the initial and subsequent driver mutations, respectively. In the simulator, we set these variants as driver mutations. Parameters were estimated based on the fit with the “observed” VAFs of the synthetic data. For real time conversion, we used a tumor size change from the diameter of 7 to 9 mm for 13 months, observed in a case of lung adenocarcinoma^16^, to convert the simulation time unit into the real time unit. The other settings are provided in **Data availability**.

The estimated parameters were close to the “true” parameters of the synthetic data (**Supplementary Figure 8a**). With estimated parameters, the tumor grew from a single cell to a clinically detectable size (10^9^ cells) in computer. As shown in **Figure 3a**, VAFs obtained from the simulated tumor fitted well with “observed” VAFs from the synthetic data (the average of the absolute mean error across the data points shown in the figure: 1.9%). The tumor content (0.8) estimated based on the least mean error was identical to the true tumor content (**Figure 3b**). Note that simulation VAFs shown in **Figure 3a** were corrected for this tumor content (**Methods**). The degrees of contribution of the driver genes to the cell-division rate increment were estimated as proportions of 0.65 and 0.35 on average for the *TP53* and *BRAF* aberrations, respectively (**Figure 3c**). The true values, 0.6 and 0.4, respectively, showed good concordance with these estimates.

**Figure 3.**
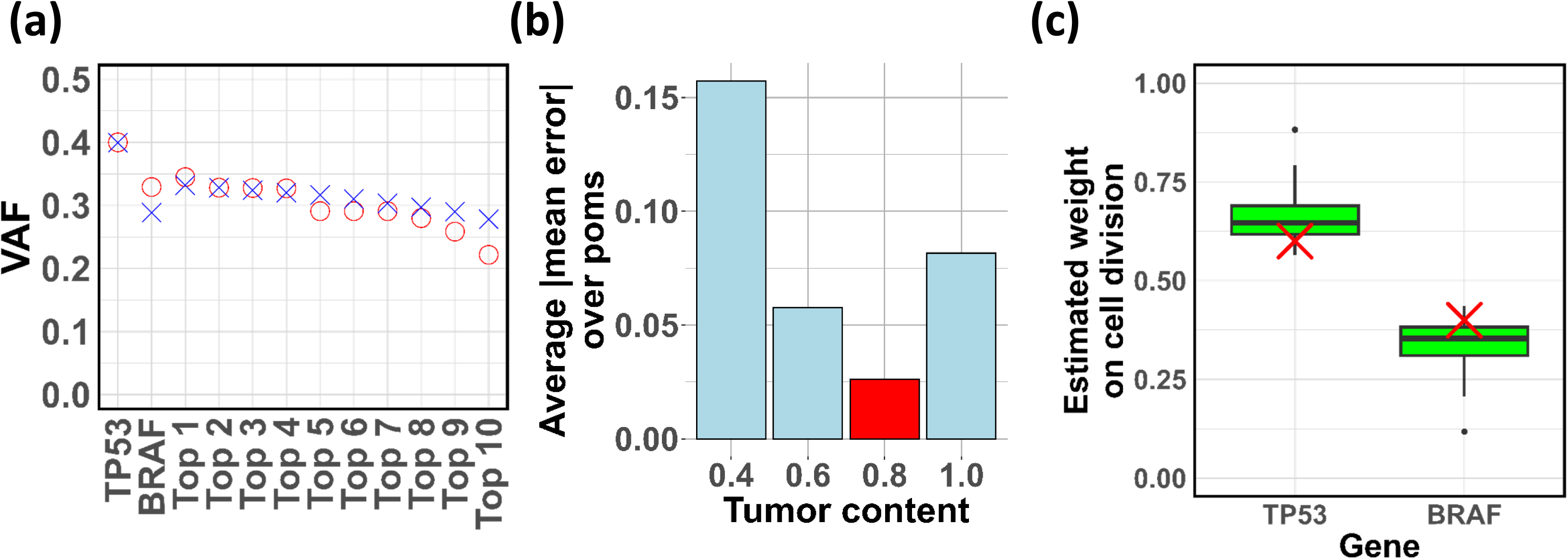
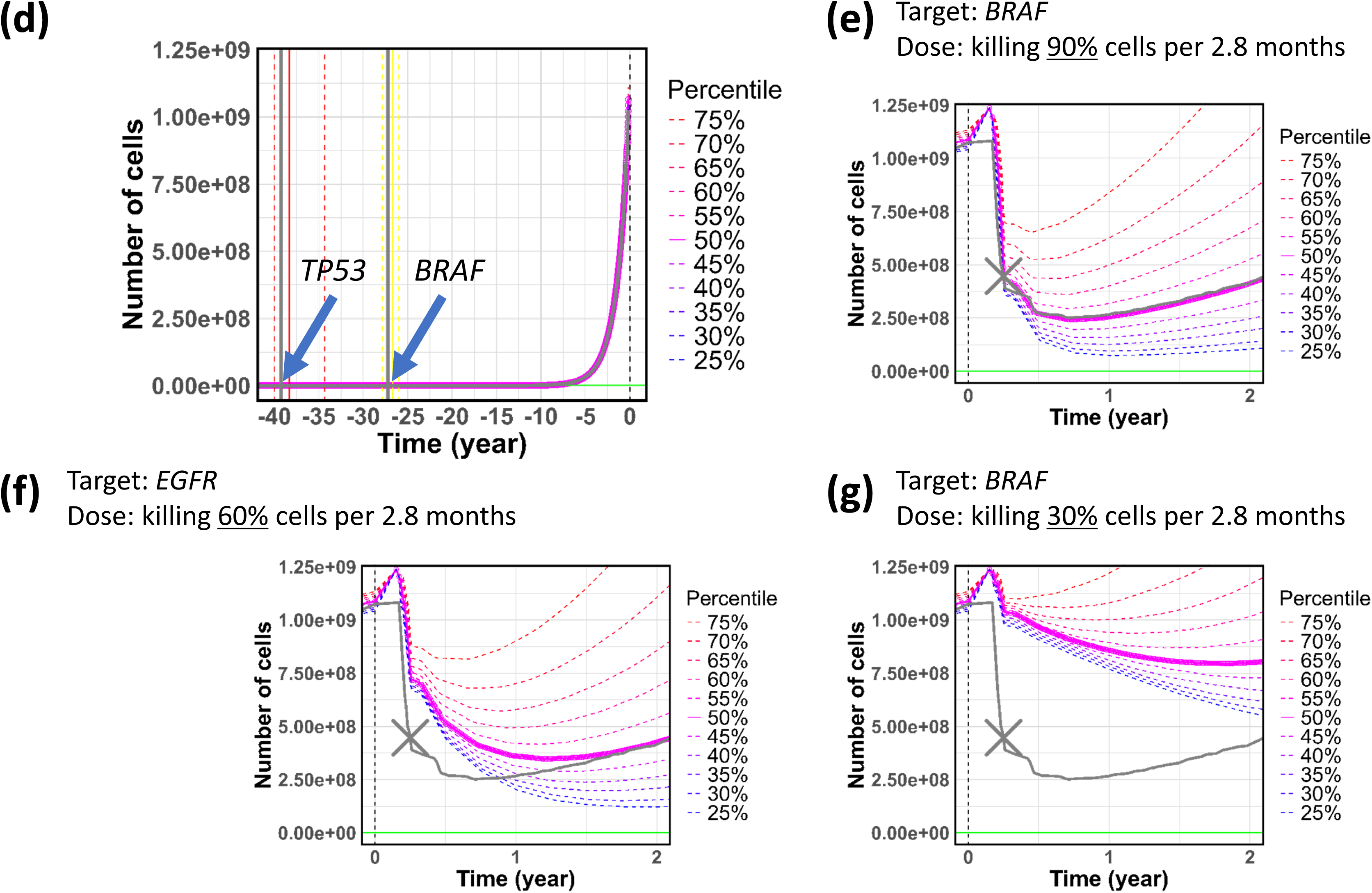
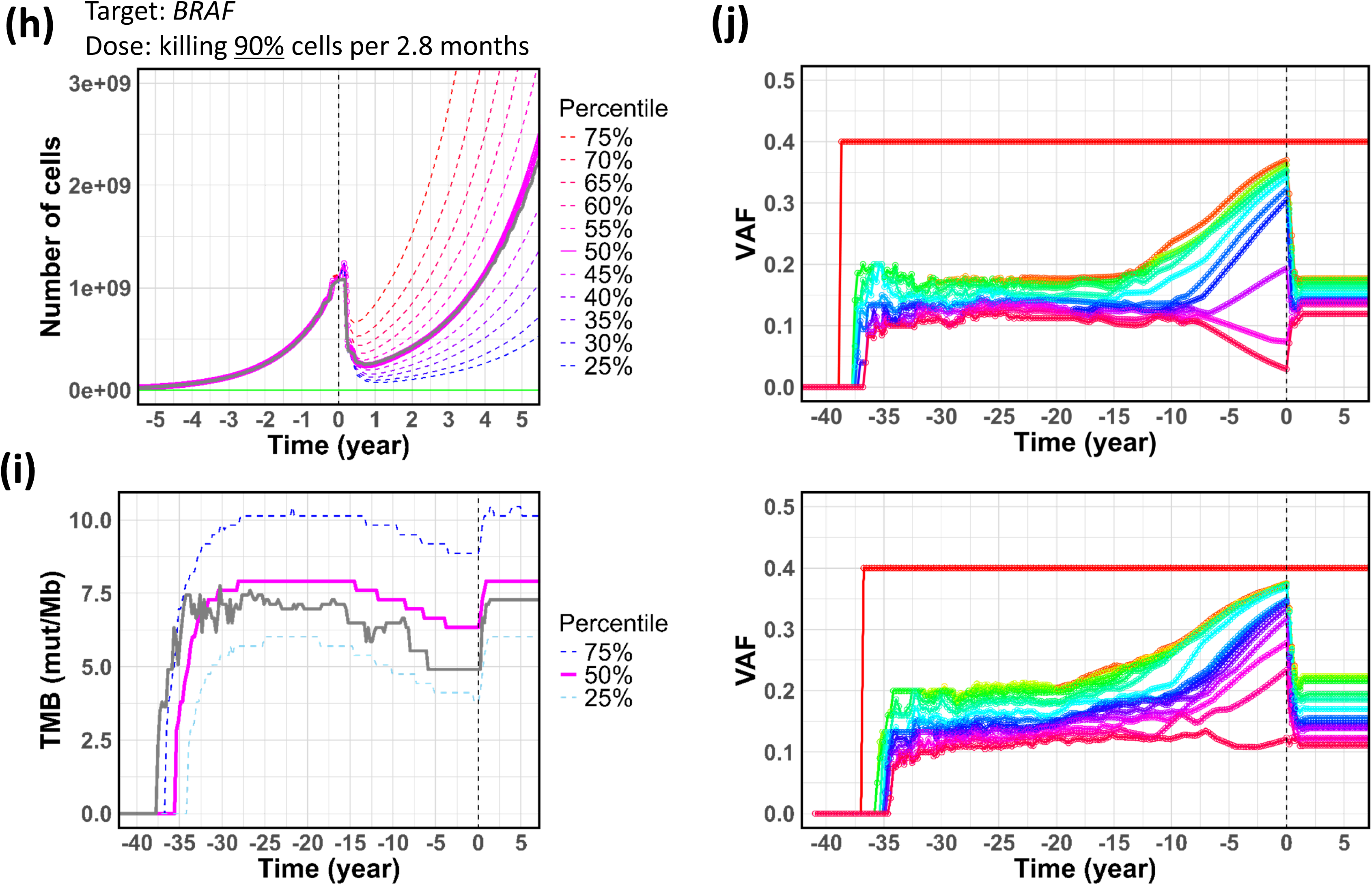
Application to synthetic data 1. **(a)** VAFs in synthetic and simulation data. The circle in red represents the “observed” VAFs from the synthetic data. The cross in blue represents the mean of the VAFs of a simulation pom across simulation replications. Top 1, 2, … represent passenger poms according to the ranks of VAFs. **(b)** Tumor content versus VAF error. For each tumor content, the absolute values of differences between observed VAFs and simulation mean VAFs were calculated and then the average was measured across poms (simulation VAFs here are values before parameter estimation by ABC). The red bar indicates the “true” tumor content in the synthetic data. **(c)** Weight parameters representing increments to the cell division rate. The cross in red represents the “true” values of the synthetic data. **(d)** Time versus number of cells. The observation and starting timepoints of drug intervention are set at time 0. The vertical lines on the left represent the timings of driver mutations (the solid, median; the dashed, 25^th^ and 75^th^ percentiles). The solid and dashed curves represent the median and other percentiles, respectively. The horizontal line in green represents a detectable level (10^6^ cells) by radiological imaging. The vertical lines and curve in gray represent the “true” ones from the synthetic data. **(e – g)** Time versus number of cells in dug intervention under different drug conditions. The cross in gray indicates the number of cells obtained from radiological imaging conducted at the corresponding time point. The curve in gray represents the “true” curve from the synthetic data. **(h)** Time versus number of cells under the killing rate. The curve in gray represents the “true” curve from the synthetic data. **(i)** Time versus TMB. The curve in gray represents the “true” curve from the synthetic data. **(j)** Time versus VAF. VAFs over 0.1 were selected and then ranked and colored according to the ranks (up to 20) at each time point. The upper shows the “true” VAFs from the synthetic data; the lower shows simulation VAFs.

The real-time conversion enabled us to estimate the timing of mutation events. The tumor was estimated to start with the *TP53* variant 38.3 (as the ensemble median across simulation replications; 34.3–40.0 as the 25^th^–75^th^ ensemble percentiles) years before reaching the clinically detectable size; the *BRAF* variant was estimated to hit 26.7 (26.0 – 27.9) years before (**Figure 3d**). The true values, 39.3 and 27.2, respectively, showed good concordance with these estimates. The simulated growth curve was almost identical to the true growth curve (**Figure 3d**).

We provided cell number curves under virtual drug intervention targeting the *BRAF* variant with doses to kill 90%, 60%, and 30% of cancer cells every 2.8 months (**Figure 3e-g**). Here, we assumed that a tumor size calculated from radiological imaging 3 months after the start of drug administration was given to estimate a corresponding killing rate (**Supplementary Figure 7b**). Because the given size was closest to the size predicted for the dose killing 90% in 3 months, the killing rate was estimated to be 90% (**Figure 3e**). Indeed, the true value was 90%. At this killing rate, the tumor was predicted to initially shrink to 1/5 of its original size in 0.7 years but regrow to the size in 3.9 years (**Figure 3h**). The predicted cell-number curve closely matched the true curve.

We estimated TMB (tumor mutation burden: the number of point mutations per Mb) and VAFs versus time. The estimated TMB dynamics was close to the true dynamics (**Figure 3i**). The overall trend of the estimated VAF dynamics was similar to the true dynamics (**Figure 3j**).

#### 2) Synthetic data 2

This synthetic dataset was generated based on a colorectal cancer patient. In this dataset, a SNV and copy-number loss in *APC*, and SNVs in *KRAS* and *TP53* were defined as driver mutations. For real time conversion, we used a tumor volume doubling time of 211 days in colon adenocarcinoma^17^, to convert the simulation time unit into the real time unit (**Methods**).

The estimated parameters closely matched the “true” parameters of the synthetic data (**Supplementary Figure 8b**). The VAFs of the simulated tumor were consistent with the “observed” VAFs of the synthetic data (the average of the absolute mean error: 2.5%) (**Figure 4a**). The estimated tumor content (1.0) was identical to the true value (**Figure 4b**). The contributions of the *APC*, *KRAS*, and *TP53* aberrations to the division rate increment was estimated as proportions of 0.50, 0.24, and 0.25 on average, respectively, which closely matched the true values of 0.50, 0.25, and 0.25 (**Figure 4c**). The three aberrations were estimated to occur 16.0 (15.6 – 16.4), 10.3 (9.9 – 10.7), and 8.9 (8.5 – 9.3) years before the observation timepoint, respectively, which also showed good agreement with the true values of 16.2, 10.6, and 9.0, respectively (**Figure 4d**). The estimated growth curve was almost identical to the true curve (**Figure 4d**).

**Figure 4.**
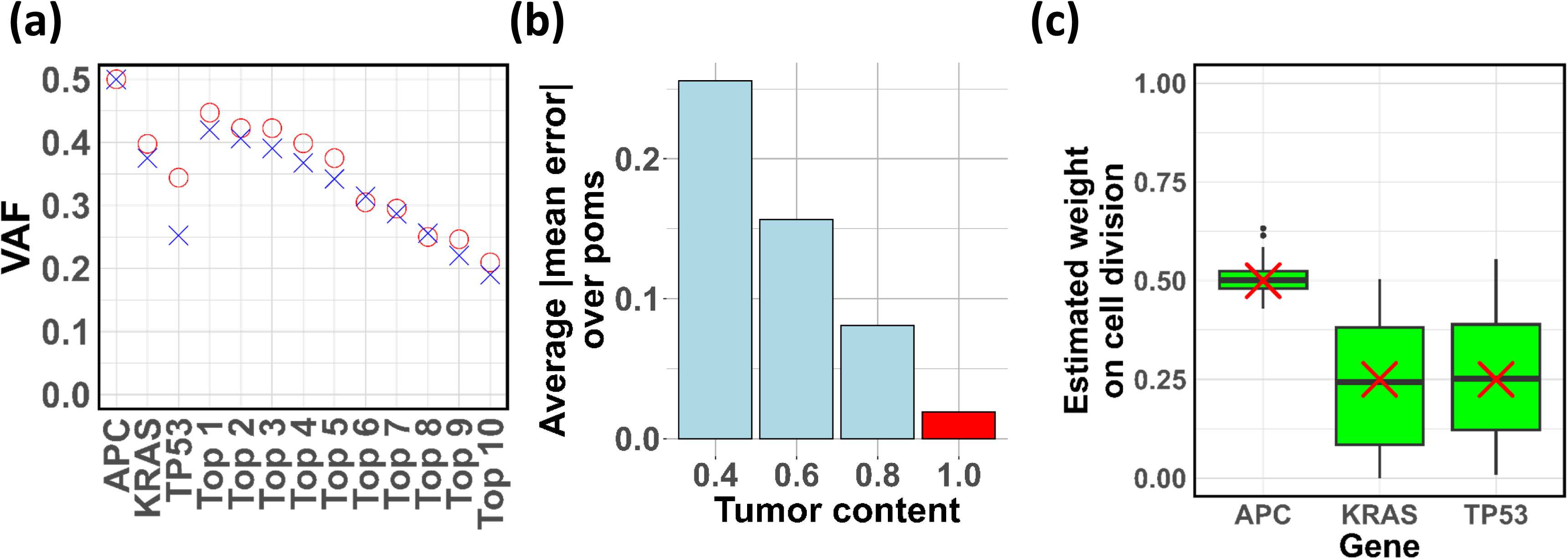
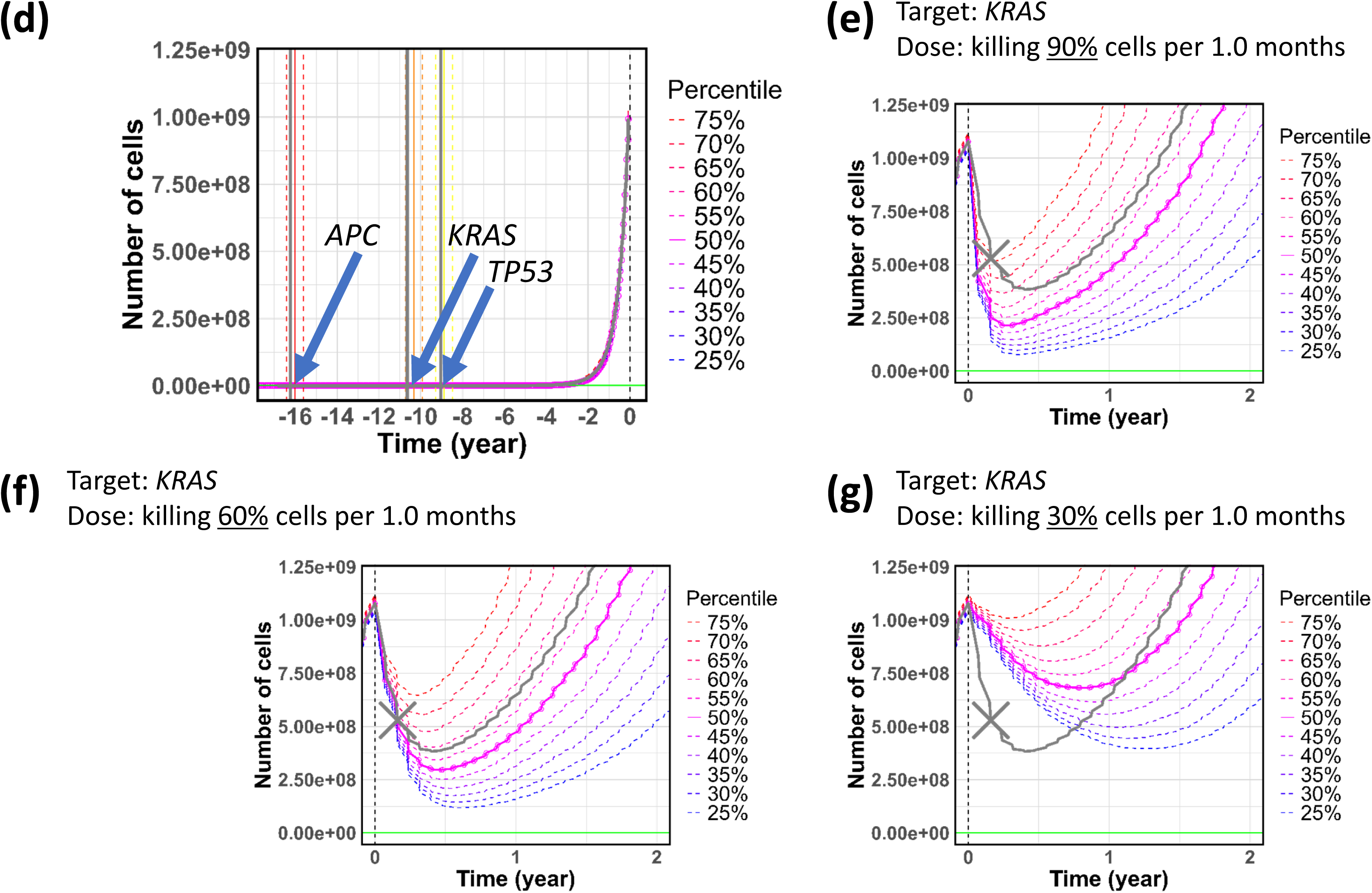
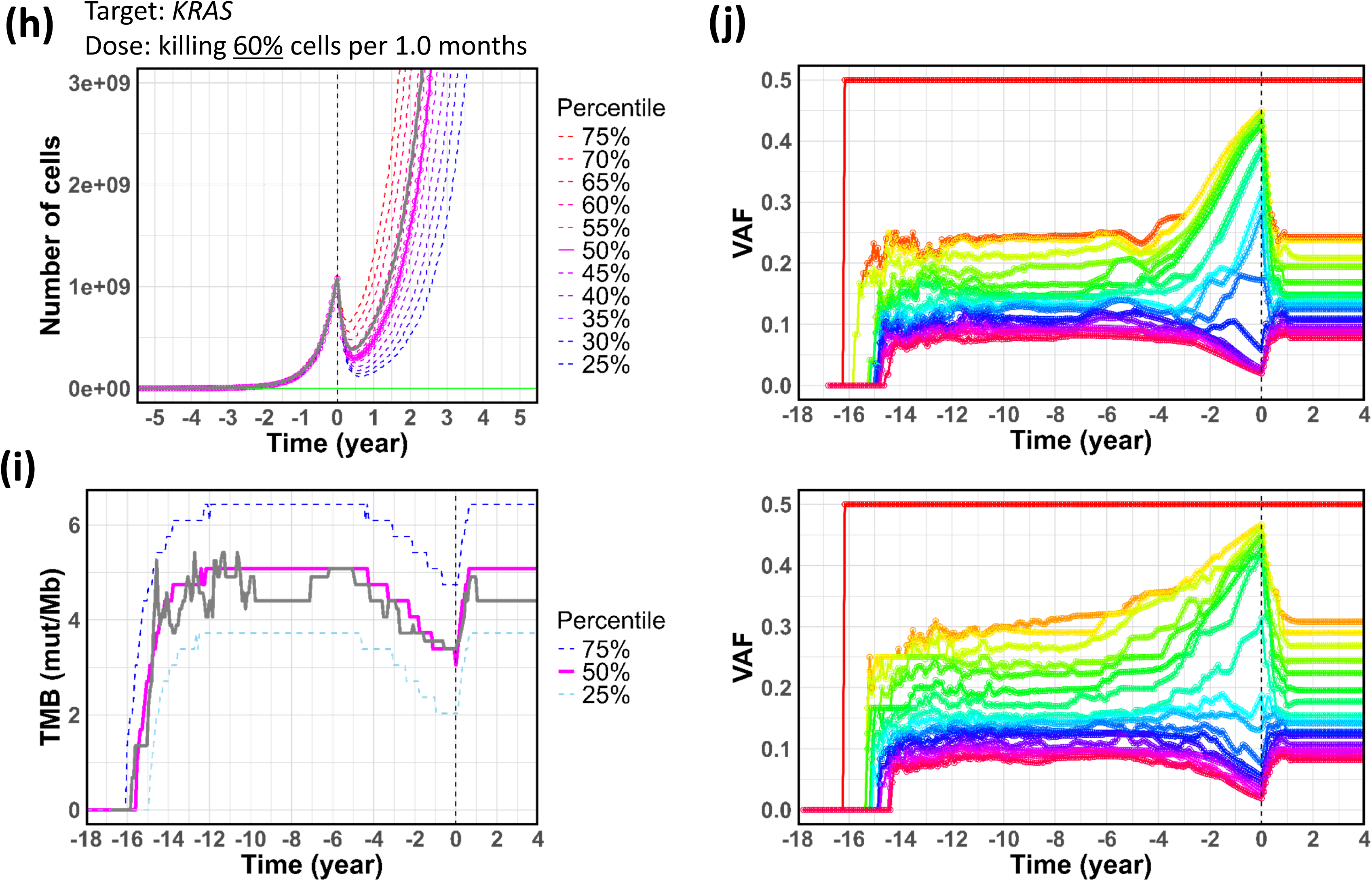
Application to synthetic data 2. **(a)** VAFs in synthetic and simulation data. The circle in red represents the “observed” VAFs from the synthetic data. The cross in blue represents the mean of the VAFs of a simulation pom across simulation replications. Top 1, 2, … represent passenger poms according to the ranks of VAFs. **(b)** Tumor content versus VAF error. For each tumor content, the absolute values of differences between observed VAFs and simulation mean VAFs were calculated and then the average was measured across poms (simulation VAFs here are values before parameter estimation by ABC). The red bar indicates the “true” tumor content in the synthetic data. **(c)** Weight parameters representing increments to the cell division rate. The cross in red represents the “true” values of the synthetic data. **(d)** Time versus number of cells. The observation and starting timepoints of drug intervention are set at time 0. The vertical lines on the left represent the timings of driver mutations (the solid, median; the dashed, 25^th^ and 75^th^ percentiles). The solid and dashed curves represent the median and other percentiles, respectively. The horizontal line in green represents a detectable level (10^6^ cells) by radiological imaging. The vertical lines and curve in gray represent the “true” ones from the synthetic data. **(e – g)** Time versus number of cells in dug intervention under different drug conditions. The cross in gray indicates the number of cells obtained from radiological imaging conducted at the corresponding time point. The curve in gray represents the “true” curve from the synthetic data. **(h)** Time versus number of cells under the killing rate. The curve in gray represents the “true” curve from the synthetic data. **(i)** Time versus TMB. The curve in gray represents the “true” curve from the synthetic data. **(j)** Time versus VAF. VAFs over 0.1 were selected and then ranked and colored according to the ranks (up to 20) at each time point. The upper shows the “true” VAFs from the synthetic data; the lower shows simulation VAFs.

We assumed that a tumor size calculated from radiological imaging 2 months after drug administration was given. The killing rate for which the predicted tumor size was closest to the given size was 60% per 1.0 month (**Figure 4e-g**). This estimated rate matched the true value of 60%. The predicted cell-number curve in the virtual drug intervention was close to the true curve (**Figure 4h**). The estimated TMB and VAF dynamics were in close agreement with the true dynamics (**Figure 4i,j**).

### Simulation examples using real data

We present simulations based on real data from patients arbitrarily selected from the TCGA database.

#### 1) TCGA 8506

The patient, referred to as 8506 (TCGA ID: TCGA-55-*8506*-01A-11D-2393-08), was a 62-year-old female with lung adenocarcinoma. In this patient, an SNV in *EGFR* and SNVs in *TP53* and *BRAF* were supposed to be the initial and subsequent driver events, respectively. In the simulator, we set these variants as driver mutations and variants in chromosome I as passenger mutations. As performed in synthetic data 1, parameters were estimated and the simulation time was converted into real time. This example is merely a demonstration of our simulator. In reality, the tumor-size change used for the time conversion should be replaced with a real change and driver genes should be determined by the tumor board of CGP tests. The other settings are provided in **Data availability**.

As shown in **Figure 5a**, VAFs obtained from simulated tumor fitted well with VAFs obtained from real TCGA data (the average of the absolute mean error across the data points shown in the figure: 2.8%), suggesting successful parameter estimation. The tumor content was estimated at 1.0 based on the least mean error (**Figure 5b**). The degrees of contribution of the driver genes to the cell-division rate increment were estimated as proportions of 0.61, 0.17, and 0.20 on average for the *EGFR*, *TP53*, and *BRAF* aberrations, respectively (**Figure 5c**).

**Figure 5.**
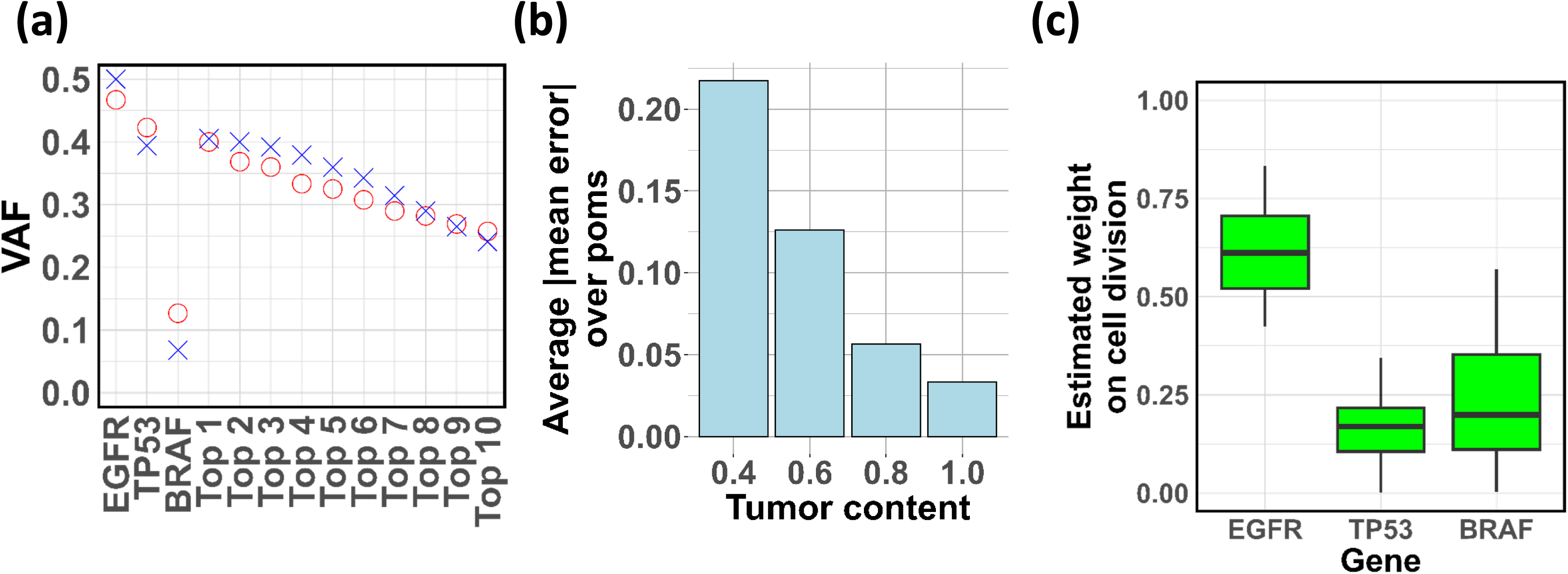
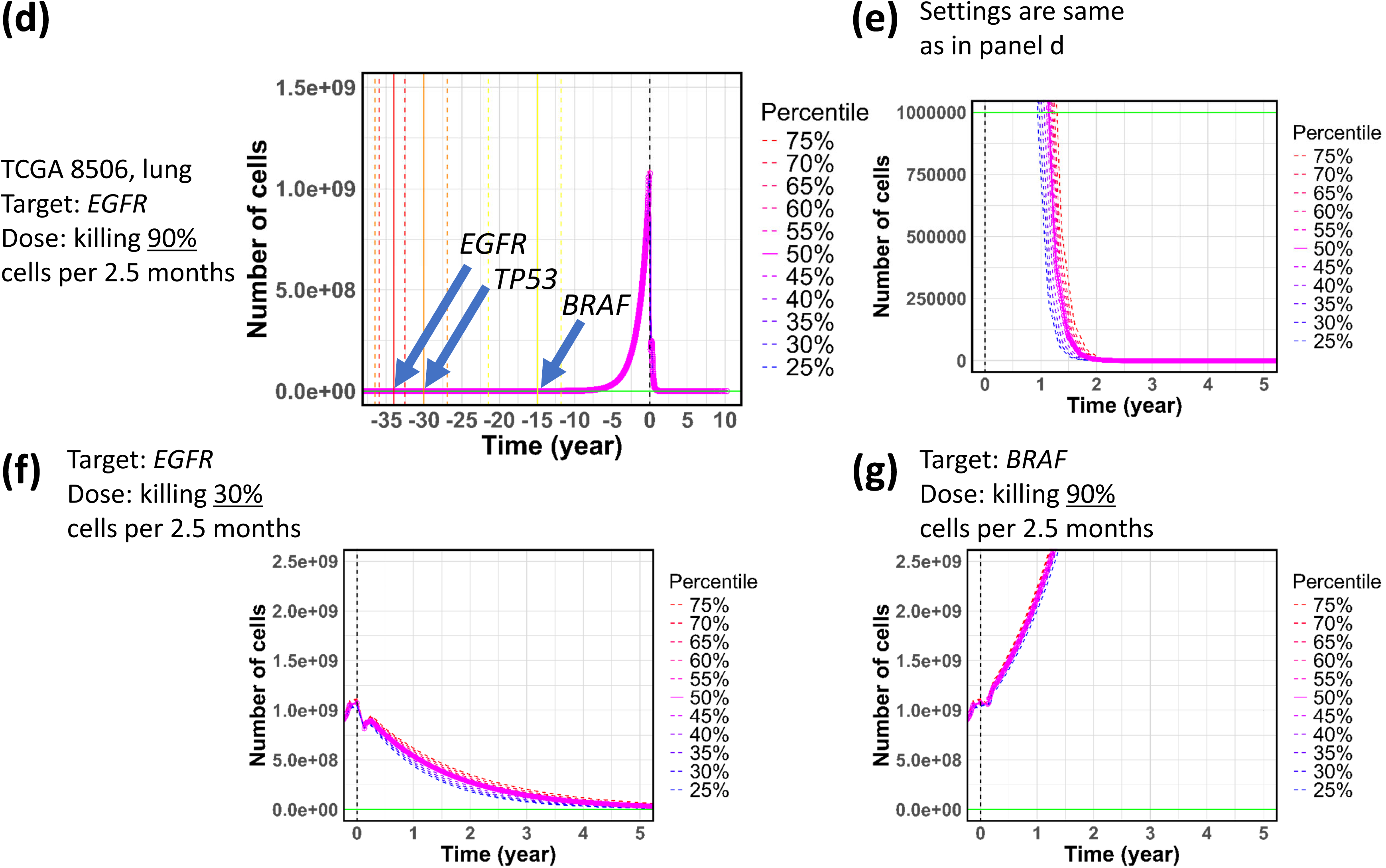
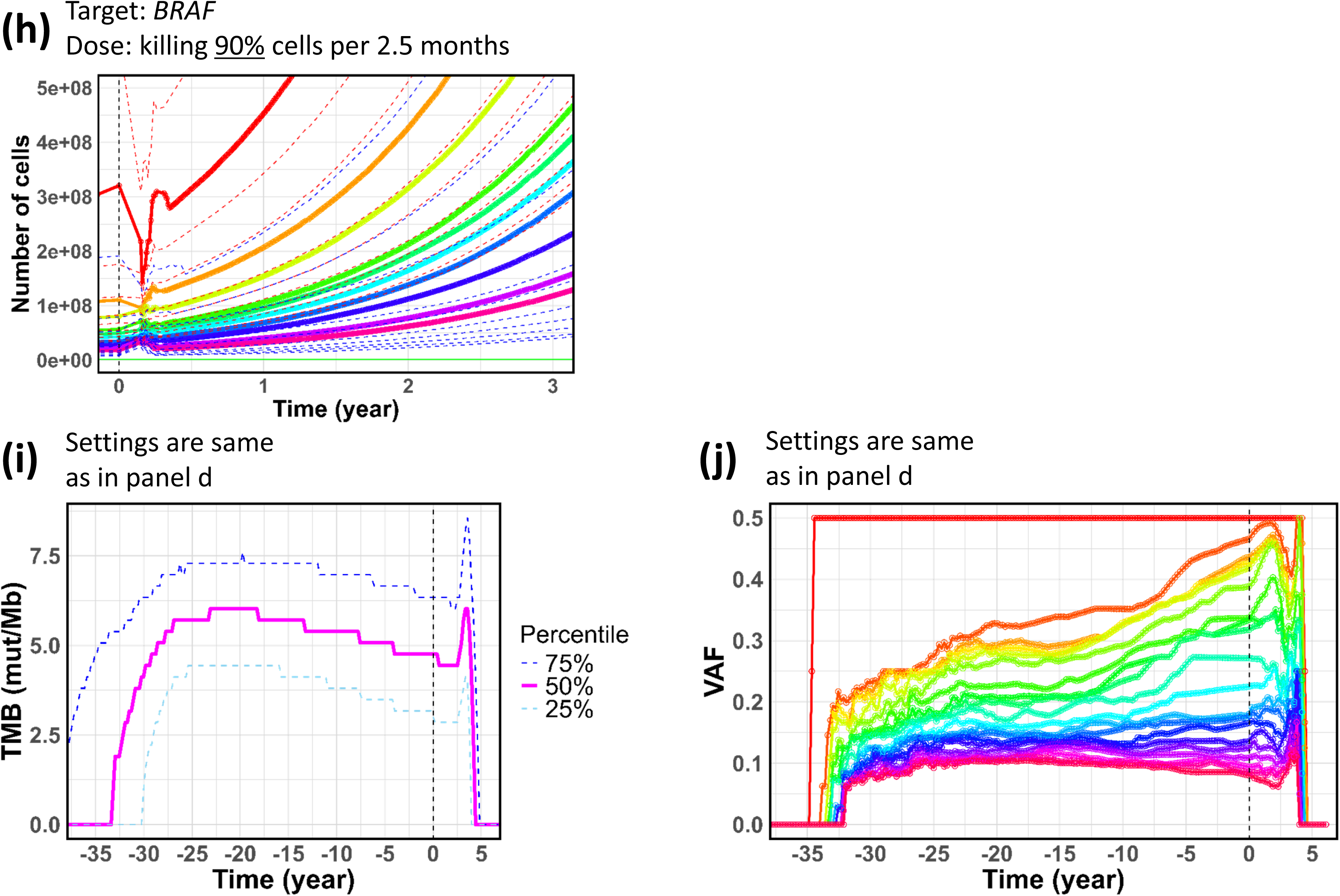
Application to TCGA 8506. **(a)** VAFs in observed and simulation data. The circle in red represents the VAFs of observed poms. The cross in blue represents the mean of the VAFs of a simulation pom across simulation replications. Top 1, 2, … represent passenger poms according to the ranks of VAFs. **(b)** Tumor content versus VAF error. For each tumor content, the absolute values of differences between observed VAFs and simulation mean VAFs were calculated and then the average was measured across poms (simulation VAFs here are values before parameter estimation by ABC). **(c)** Estimated weight parameters representing increments to the cell division rate. **(d)** Time versus number of cells. The observation and starting timepoints of drug intervention are set at time 0. The vertical lines on the left represent the timings of driver mutations (the solid, median; the dashed, 25^th^ and 75^th^ percentiles). The solid and dashed curves represent the median and other percentiles, respectively. The horizontal line in green represents a detectable level (10^6^ cells) by radiological imaging. **(e – g)** Time versus number of cells in dug intervention under different drug conditions. **(h)** Time versus number of cells for subclones. The curves in different colors represent the top 10 subclones in the number of cells at each time point. The horizontal line in green represents the detectable level. The dashed curves in blue and red represent 25^th^ and 75^th^ percentiles of each subclone. **(i)** Time versus TMB. **(j)** Time versus VAF. VAFs over 0.1 were selected and then ranked and colored according to the ranks (up to 20) at each time point.

The tumor was estimated to start with the *EGFR* variant 34.0 (as the ensemble median across simulation replications; 32.5–36.0 as the 25^th^–75^th^ ensemble percentiles) years before reaching the clinically detectable size. The subsequent *TP53* and *BRAF* variants were estimated to hit 30.0 (26.9 – 36.5) and 14.9 (11.8 – 21.4) years before, respectively (**Figure 5d**).

In virtual drug intervention, we provided the computationally grown tumor with virtual drug targeting the *EGFR* variant with a dose to kill 90% of cancer cells every 2.5 months. The ensemble prediction showed that the tumor would shrink below a visible level (10^6^ cells) on the radiological image in 1.2 years and be eliminated in 4.3 years on average (based on the ensemble median) (**Figure 5e**). When we used a drug with a lesser dose, corresponding to achieving a killing effect of 30% of cancer cells per 2.5 months, it would take more than 10 years for cancer cells to be invisible (**Figure 5f**).

When we targeted *BRAF* with a killing rate of 90%, the targeted therapy would fail; the tumor would rapidly grow after a brief pause (**Figure 5g**). This regrowth came from the presence of clones without the targeted *BRAF* variant. When the drug was administered, major clones somewhat decreased until 0.2 years; however, clones without the *BRAF* variant steadily increased and overtook previous major clones (**Figure 5h**).

We made an ensemble prediction of TMB and VAFs versus time. We excluded point mutations with VAFs of less than a threshold (conservative, 10%) from the calculation of TMB, as done in reality. Real TMB was observed at time 0 (observation timepoint) and the value was 7.5 (decreased to 1 / 20 of the original for convenience, see **Methods**), while TMB in the simulation average was 4.8 (3.0 – 6.3) (**Figure 5i**).

TMB drastically increased at the onset, rising from 0 to 6.0, and then gradually decreased until the observation timepoint (**Figure 5i**). After the drug intervention (killing rate of 90%), TMB would slowly decline before suddenly increasing, peaking in 4 years, during which the tumor size would drastically decrease. This increase in TMB may seem counter-intuitive but it occurred because minor clones with poms under the VAF threshold relatively increased as the drug eliminated major clones, ultimately causing the VAFs of poms of minor clones to rise beyond the threshold. Finally, TMB would drop down to zero rapidly. VAFs initially rose rapidly at the onset and then higher VAFs slowly increased until the observation timepoint (**Figure 5j**). After the drug intervention, the diverged values appeared to converge around 0.25.

#### 2) TCGA DM

The last example is of a 72-year-old female with colon adenocarcinoma, referred to as DM (TCGA ID: TCGA-*DM*-A28E-01A-11D-A16V-10). In this patient, SNVs in *APC*, *TP53*, and *KRAS*, and a short deletion in *APC* were considered driver mutations. Real time conversion was performed as in synthetic data 2. We set the simulator and tuned the parameters as in TCGA 8506.

The observed VAFs were well reproduced in our simulator (the average of the absolute mean error: 4.8%) with an estimated tumor content of 1.0 (**Figure 6a,b**). Notably, LOH in *TP53* (the variant with observed and simulated VAFs of 0.8 and 0.71, respectively, in **Figure 6a**) was reproduced. The contributions of *APC*, *KRAS*, and *TP53* to the division rate increment were estimated as proportions of 0.26, 0.21, and 0.54 on average, respectively (**Figure 6c**). The tumor was estimated to initiate 16.4 (15.6 – 16.8) years before the observation timepoint, and the *KRAS* mutation, *TP53* loss, and *APC* deletion were estimated to occur 12.2 (11.3 – 12.6), 10.9 (10.1 – 11.3), and 9.9 (9.1 – 10.4) years before, respectively (**Figure 6d**).

**Figure 6.**
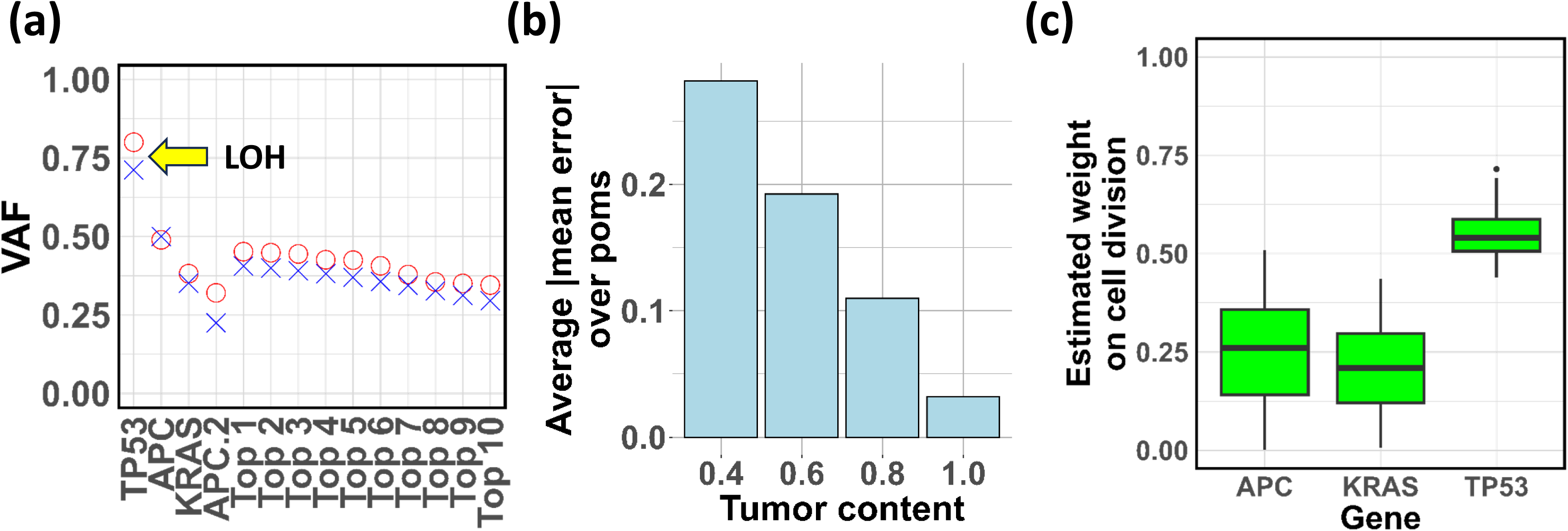
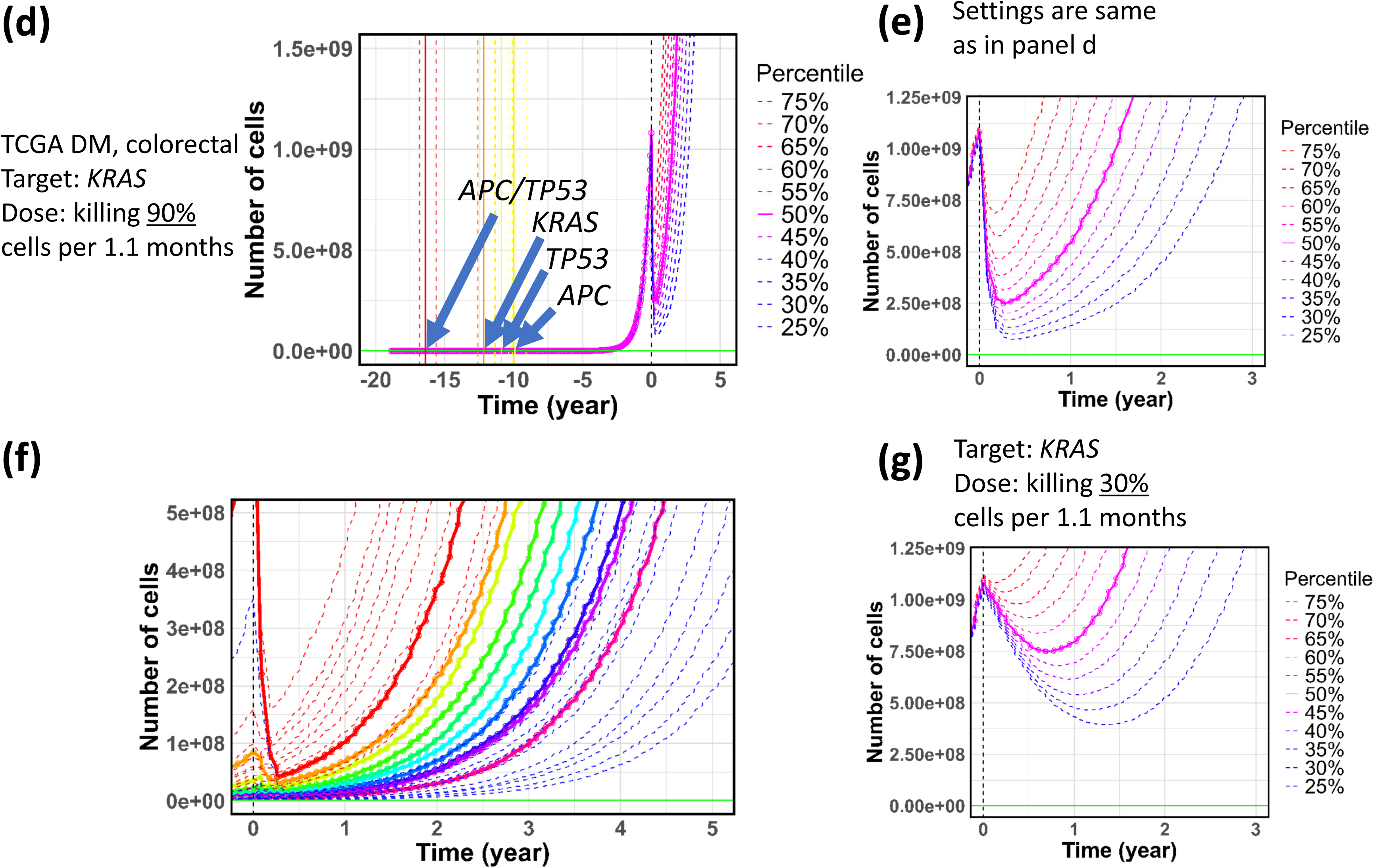
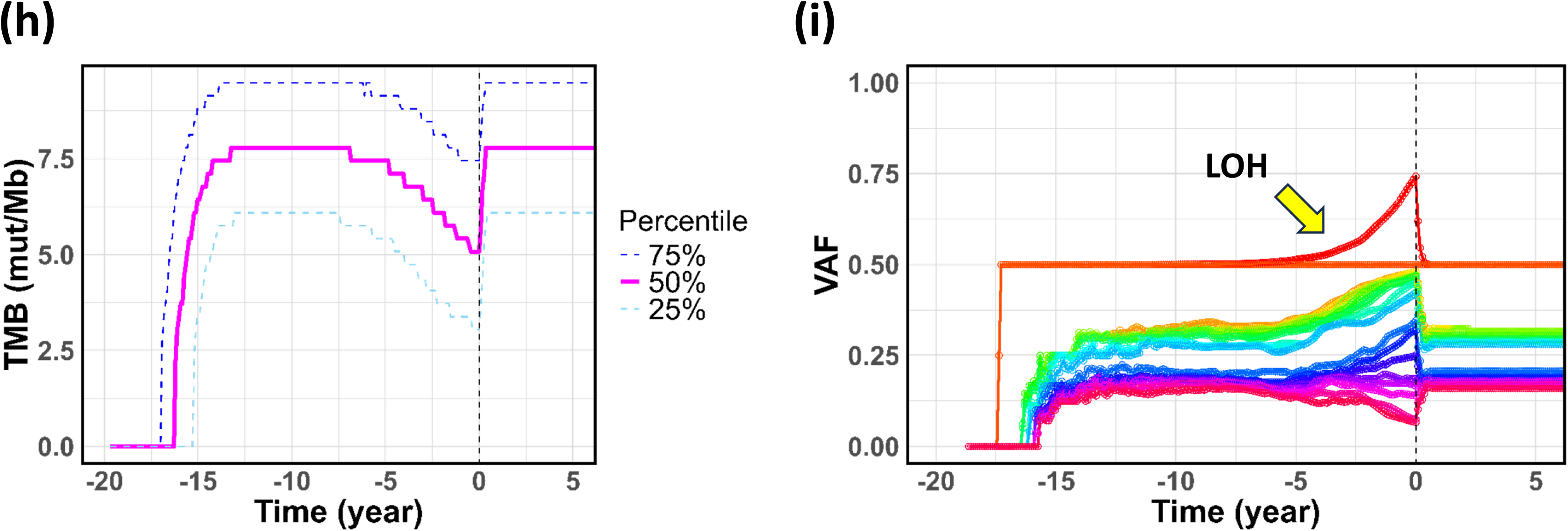
Application to TCGA DM. **(a)** VAFs in observed and simulation data. The circle in red represents the VAFs of observed poms. The cross in blue represents the mean of the VAFs of a simulation pom across simulation replications. Top 1, 2, … represent passenger poms according to the ranks of VAFs. **(b)** Tumor content versus VAF error. For each tumor content, the absolute values of differences between observed VAFs and simulation mean VAFs were calculated and then the average was measured across poms (simulation VAFs here are values before parameter estimation by ABC). **(c)** Estimated weight parameters representing increments to the cell division rate. **(d)** Time versus number of cells. The observation and starting timepoints of drug intervention are set at time 0. The vertical lines on the left represent the timings of driver mutations (the solid, median; the dashed, 25^th^ and 75^th^ percentiles). The solid and dashed curves represent the median and other percentiles, respectively. The horizontal line in green represents a detectable level (10^6^ cells) by radiological imaging. **(e, g)** Time versus number of cells in dug intervention under different drug conditions. **(f)** Time versus number of cells for subclones. The curves in different colors represent the top 10 subclones in the number of cells at each time point. The horizontal line in green represents the detectable level. The dashed curves in blue and red represent 25^th^ and 75^th^ percentiles of each subclone. **(h)** Time versus TMB. **(i)** Time versus VAF. VAFs over 0.1 were selected and then ranked and colored according to the ranks (up to 20) at each time point.

Targeting *KRAS* with a dose equivalent to killing 90% of cells per 1.1 months would result in a sharp decrease in cancer cells until 0.3 years on average (**Figure 6e)**. However, the tumor would regrow and return to its original size in 1.6 years. This regrowth was due to the presence of clones without the *KRAS* aberration. The most dominant clone would rapidly decrease to 1/20 of its original size in 0.3 years; however, clones without the aberration would persist and displace previously major clones (**Figure 6f**). A reduced killing rate of 30% would only decrease the tumor to 70% of its original size, and it would regrow to the size in 1.5 years (**Figure 6g**).

The real and simulation TMB at the observation timepoint were 4.1 and 5.1 (3.0 – 7.5), respectively, showing a reasonable concordance (**Figure 6h**). After drug intervention with a dose corresponding to a killing rate of 90% per 1.1 months, TMB would drastically increase to 7.8 mut/Mb in 0.3 years, during which the tumor would drastically decrease. In the VAF dynamics, the LOH mutation (VAF > 0.5) appeared 11.2 years before the observation timepoint and the VAF steadily increased until the drug intervention (**Figure 6i**). After the intervention, the LOH VAF would sharply decrease because of the cells being killed and then become indistinguishable from other VAFs.

## DISCUSSION

We developed a realistic cancer-evolution simulator for personalized medicine and demonstrated its application to targeted therapies in cancer genome medicine. This demonstration is based on the exome sequencing of TCGA but is easily extendable to target sequencing performed in CGP tests. Required information is: 1) VAF data, routinely and subsidiarily obtained with NGS in cancer-genomic analyses and CGP tests, respectively; 2) radiological images at two time points for the real time conversion, which can be routinely acquired in monitoring cancer; and 3) driver genes, routinely determined by the tumor board in CGP tests for drug targets. To calibrate drug killing rate, 4) radiological image at one time point after drug administration is needed. All these items are personalized data for personalized medicine, though average data can be used if personalized data are unavailable.

Predictions made by the simulator can guide treatment plans. Our demonstration exemplified targeted therapies that would fail because targets or the dosage was inappropriate despite the therapies targeting driver genes. It also showed the tentative surge in TMB despite drug response, calling for caution against unnecessary immune checkpoint therapies. Such predictions would be especially valuable when cancer cells decrease below the detection limit in radiological images after treatment. Simulations may also indicate an appropriate timepoint for liquid biopsy to monitor cancer-cell levels (*e.g.*, for minimum residual disease), further combined with data assimilation such as Kalman filtering for better planning.

Previous genome-oriented simulators of cancer evolution^4–15,18–21^ focused on basic biology, such as understanding ITH, and were not designed for realistic applications in targeted therapy within cancer genome medicine. While another group of cancer simulators based on non-genomic data^22–26^ exists, their starting points differ, leaving the utilization of genomic data obtained through genome medicine, such as CGP tests, unclear.

We used basic ABC and BO techniques with limited summary statistics for parameter estimation. Advanced techniques and additional summary statistics could yield more precise predictions. We did not model spatial information because current CGP tests lack it. Nevertheless, recent studies^11,18,27–29^ suggest that incorporating spatial structures could better capture ITH, potentially improving our model. We plan to validate predictions from virtual drug interventions using genome-sequenced patient-derived xenografts, though this is beyond the scope of this study focusing on theoretical and computational aspects. Drug-resistant cells and gene-gene interactions were not modeled and remain future research directions. tugMedi was developed for research purposes and is not intended for clinical use.

Although the simulation model in this study is only a first approximation of the real tumor, we demonstrated the application of the simulator to targeted therapies in cancer genome medicine. This study is the first step toward the numerical prediction of cancer care, similar to numerical weather prediction for forecasting the course of typhoons to help reduce disaster impacts.

## METHODS

### CNA

We introduced CNAs in our model. Changes in simulation processes related to CNAs from previous studies^14,21^ are overviewed in **Supplementary Figure 2a,b**. The concept of parental chromosomes is introduced, and that of mutation is divided into point mutation (abbreviated as pom), deletion (del), and duplication (dup) (**Supplementary Figure 2a**). Poms are SNVs and indels in reality. Dels and dups are the main causes of CNAs^30–32^. Each mutation type can occur on a parental chromosome and cause gene malfunction given the malfunction probability of each mutation type. Unlike poms, dels and dups (CNAs) have a length feature and may cause malfunction in multiple genes in a single event.

A pom occurs only on the exons of genes, for which the exon-intron structures are input by users. Then, a pom generates a variant allele B, different from the original allele A, which is located on the other parental chromosome (**Supplementary Figure 2b**). We do not suppose more than one mutation at one site due to an extremely low occurrence probability (only variant B, no other variants C, D, or others).

We modeled the mechanisms of dels and dups, in which a parental chromosome is shrunk or extended by a chromosomal segment between the two breakpoints of a del or dup, respectively (**Supplementary Figure 2c**). The deletion-shrink model has been proposed previously^33^. The reciprocal mechanism of del and dup events is often observed^31,32^ and is represented in this study by the combination of the deletion-shrink and duplication-extension models (**Supplementary Figure 2c**). According to shrink and extension, the lengths of chromosomes are changed and thereby differ between parental chromosomes and between clones.

For a del or dup event, the first breakpoint stochastically occurs following a Poisson distribution with the mean of the exon + intron length multiplied by the occurrence probability of del or dup, respectively (**Supplementary Figure 2d**). Given its Poisson distribution, multiple first breakpoints may occur at one time. The chromosomal position of a first breakpoint is stochastically determined following a uniform distribution among possible chromosomal positions. After this “fall” process of a first breakpoint, the paired second breakpoint is determined by “extension” process, where the length of a chromosomal segment for a del or dup is stochastically determined following an exponential distribution with a del/dup average length provided by users (**Supplementary Figure 2d**). Deletion lengths may follow either power-law or exponential distribution but are practically well-approximated by an exponential distribution within an effective range^33^. We adopted an exponential distribution for its simpler calculation. More details on the implementation of CNA are described in **Supplementary Methods** and illustrated in **Supplementary Figure 3**.

Genes malfunction according to the malfunction probability of each mutation type (**Supplementary Figure 2e**). A point mutation may result in the malfunction of a single gene in one event. A deletion or duplication may result in the malfunction of multiple genes covered by the CNA segment individually in one event.

### VAF for given CNA and tumor purity

Considering CNA and tumor purity, we precisely defined VAF in this study. To this end, we sorted several terms. We defined a *clone* or *subpopulation* as a group of cells with identical mutations (poms and CNAs), irrespective of driver or passenger mutations. *Driver mutations* are those influencing the cell division rate (generally, influencing hallmark variables in **Supplementary Methods**), and stochastically influence cell states at the next time step^14^. Thus, driver mutations can influence the cell phenotype of the division rate^14^. *Passenger mutations* do not influence the division rate or cell phenotype.

We defined *tumor* or *cancer cells* as cells with one or more driver mutations, and *normal cells* as those without any driver mutations. However, normal cells can have passenger mutations. We defined *speckled* normal cells as normal cells with passenger mutations, and *intact* normal cells as those without any passenger mutations. Tumor purity/content is the fraction of tumor cells in a sampled specimen. The purity/content of speckled normal cells is the fraction of speckled normal cells in a sampled specimen (**Supplementary Figure 5a**).

Integrated and extended from equations in the literature – *b_i_* in ^34^; *h_f_* in ^35^; *vaf* in ^36^, a generalized form of VAF including multiple cell subpopulations in a sampled specimen is as follows:

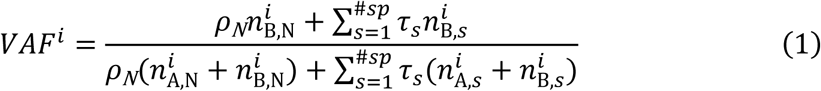

where *i* is the index for the variant position (site), *ρ*_N_ is the admixture rate of intact normal cells in a sampled specimen, *n* is the copy number of original allele A or variant B, N indicates *intact* normal cells, and *s* indicates the index for a subpopulation of tumor and *speckled normal* cells (**Supplementary Figure 5a**). *#sp* is the number of subpopulations. τ is the fraction of a subpopulation in a sampled specimen; therefore,

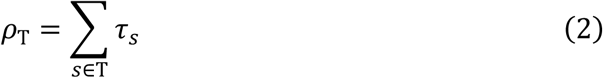

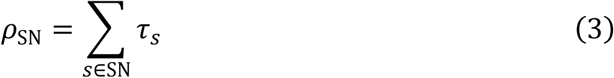

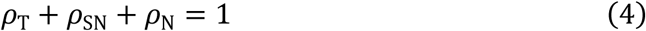

where *ρ*_T_ and *ρ*_SN_ are the tumor and speckled normal contents in a sampled specimen, respectively (**Supplementary Figure 5a**).

Tumor purity pathologically measured is usually based on the identification of cancer cells through immunostaining and cell shapes and sizes; hence, it can correspond to *ρ*_T_ (**Supplementary Figure 5a**). Because tumor cells are usually not distinguished from speckled normal cells in NGS fastq data, tumor purity estimated using SNVs in NGS data^37,38^ may include a speckled normal purity, *i.e.*, *ρ*_T_+ *ρ*_SN_ (**Supplementary Figure 5a**). If speckled normal cells exist, the tumor purity pathologically measured is theoretically smaller than the NGS-based tumor purity by *ρ*_SN_. In typical human cases, 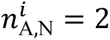 and 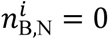 for intact normal cells. Therefore, the full equation for VAF above is reduced into a simplified form:

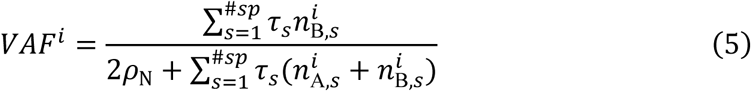

*ρ*_N_ can be estimated based on the fit between observed VAFs and simulation VAFs calculated with this equation. The method of estimation is described in **Supplementary Methods**.

### Gene structures

Poms occur in exons but CNAs occur irrespective of exons and introns in tugMedi. To define exon and intron structures, we used real gene structures in the NCBI Consensus CDS database^39^.

### Genetic modes

We introduced recessive and dominant modes in tugMedi. In the recessive mode, the malfunction states of a gene in *both* parental chromosomes are required for the genuine malfunction state of a gene, which changes the cell division rate (generally, changes hallmark variables in **Supplementary Methods**) (**Supplementary Figure 4a**). In the dominant mode, a malfunctional state in *either* parental chromosome is sufficient for the genuine malfunction state (**Supplementary Figure 4a**).

In our model, poms and dups in oncogenes follow the dominant mode, and dels in oncogenes do not exert any effects on cell phenotypes (**Supplementary Figure 4b**). Poms and dels in suppressors follow the recessive mode, and dups in suppressors do not exert any effects (**Supplementary Figure 4b**). The malfunction rate of oncogenes is defined separately from suppressors, where the former rate is usually lower than the latter because of the gain-of-function property in contrast to the loss-of-function property^40^.

Dominant-negative genes resemble suppressors, and the only difference is that poms in dominant-negative genes follow the dominant, not the recessive mode, as expected in *TP53* (**Supplementary Figure 4b**). Users (or generators randomly) select a type of oncogene, suppressor, or dominant-negative for each gene in regions of interest.

### Event enforcement & Fitting evaluation (EF) algorithm

It is rare to see in simulations that mutations occur within a few specific genes under biologically realistic parameter values. Many simulations are required for the occurrence. The EF algorithm speeds it up, enforcing the occurrence of such rare events. To this end, we first divided regions where mutations occur into 1) regions of interest and 2) other regions. Regions of interest are distinguishable regions where driver events have occurred, corresponding to a few specific genes, such as *APC*, *KRAS* and *TP53* in a patient with colorectal cancer. Passenger mutations can also occur in regions of interest. In contrast, other regions are indistinguishable, such as non-cancer-related genes, where only passenger mutations occur. Regions of interest generally own a smaller size than other regions. Statistics on simulated mutations in regions of interest and other regions are both used for parameter estimation.

#### EF algorithm for regions of interest

Before starting a simulation, users need to specify driver mutations, such as c.2573T>G causing p.L858R, in regions of interest, such as in *EGFR*. Specifying passenger mutations in regions of interest is optional. Typically, one to three driver mutations and no passenger mutations are specified. Users also need to specify a prior distribution to generate simulation time steps at which such specific mutations are enforced to occur in a simulation (**Supplementary Figure 1a**). As a prior, a uniform or normal distribution left-truncated at zero is typically used to generate positive values, using the rtrunc function of the truncdist library in R. Generated values are rounded to positive integers to represent simulation time steps.

Subsequently, a simulation starts, and the specific mutations are inserted into a single cell at the generated simulation time steps. Single cells into which mutations are inserted are randomly selected if users do not restrict clones where mutations occur. Users can restrict, by specifying other mutations owned by clones, such that mutations occur only in clones with the specified mutations (**Supplementary Figure 1a**). For example, when users specify mutation M1 for mutation M2, M2 will occur only in clones with M1 at the generated simulation time step (an error is output if there are no clones with M1 at the time step). This restriction is critical for efficiently simulating multi-stage tumorigenesis. The validity of the generated simulation time steps and selected cells/clones are ultimately evaluated based on the fit of summary statistics in simulated data with those in observed data.

#### EF algorithm for other regions

EF algorithm for other regions is based on a Poisson process. A timing when a mutational event is enforced to occur is sampled from an exponential distribution with a Poisson parameter. The Poisson parameter for pom and CNA is:

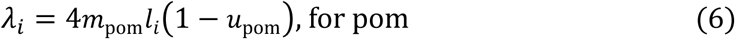

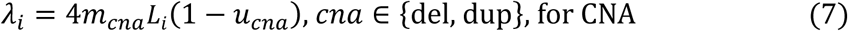

where *m*_pom_, *l*_i_, and *u*_pom_ are the pom rate per bp per cell division, the exon size of region *i* in bp, and the gene malfunction rate per pom per cell division, respectively. A factor of 4 is the correction for two parental chromosomes times two divided cells. *m*_cna_, *L*_i_, and *u*_cna_ are the occurrence rate of a CNA (del or dup) per bp per division, the exon + intron size of region *i* in bp, and the gene malfunction rate per CNA per division, respectively.

Specifically, before a simulation, 1) users need to specify the number of passenger mutations to occur. 2) For each mutation, either pom, del, or dup is randomly determined according to the relative probabilities:

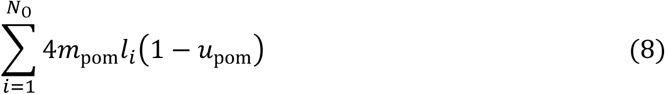

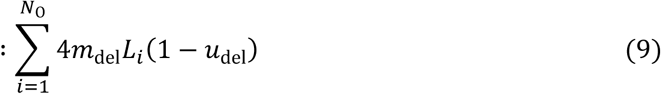

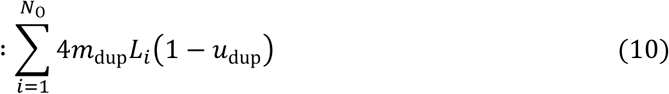

where *N*_O_ is the number of other regions. 3) Region *i* is randomly determined according to the relative probabilities:

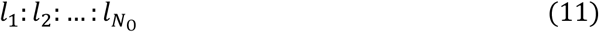

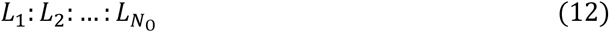

for a pom and CNA (del or dup), respectively. **Supplementary Figure 1b** illustrates three mutations randomly selected as a pom, a dup, and a pom, of which the regions are then randomly selected as regions 2, 2, and 1, respectively.

4) A random number is sampled from an exponential distribution with the rate of the Poisson parameter above (*i.e.*, with the inverse of the Poisson mean). Because the mutation rate is defined *per cell division*, this random number indicates the number of cell divisions required until a mutational event occurs. In a Poisson process, a random number drawn from an exponential distribution with a rate *per time* is referred to as *waiting time*. Accordingly, we refer to the random number as *waiting division*. **Supplementary Figure 1b** illustrates waiting divisions generated as 2, 3, and 7.

5) A simulation begins. Each mutation is inserted into a randomly selected region at the cell division timing specified by a waiting division. **Supplementary Figure 1b** illustrates the three mutations inserted into regions 2, 2, and 1 at 2^nd^, 3^rd^, and 7^th^ cell divisions, respectively.

6) For a pom, a site at the bp level, a parental chromosome, and a divided cell where a pom is inserted are randomly determined. For a CNA, the length is randomly determined from an exponential distribution as described above in the subsection on CNA. Next, the position of a CNA is uniform-randomly determined so that it will overlap with region *i* (**Supplementary Figure 1c**). A divided cell and a parental chromosome where a CNA is inserted are randomly determined.

### Conversion of simulation time into real time

Let us suppose that we obtain a real time interval in which a tumor grows from *N*_1_ cells at the first real observation to *N*_2_ cells at the second observation (**Supplementary Figure 7a**). *N*_1_ and *N*_2_ are calculated from tumor diameters obtained from radiological images through a conversion rate, such as a diameter equivalent to volume 1 mm^3^ corresponding 10^6^ cells (10^6^ cells / mm^3^)^41^. We use month as the real time unit for convenience, though it can be replaced with year or day. Meanwhile, let us suppose that we find two simulation time steps between which a *simulated* tumor grows *N*_1_ to *N*_2_ cells (**Supplementary Figure 7a**).

When we denote the real and simulation time intervals with Δ*T*(*N*_1_ → *N*_2_) and Δ*t*(*N*_1_ → *N*_2_), respectively,

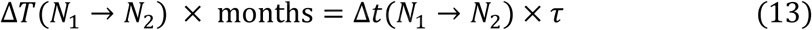

where r is the simulation time unit. Therefore, the simulation time unit is associated with the real time:

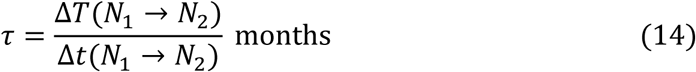

Instead of imaging observations, the classical theory of tumor volume doubling time (VDT)^41^ can be used for a rough estimation. Suppose we find a s*imulation* time interval in which a tumor roughly shows an exponential growth from *N*_1_ to *N*_2_ cells, regardless of the primary or metastatic tumor type. Suppose we know *VDT* in a real time unit of month for the primary or metastatic tumor type. When tumor types are matched between the simulation and real, the classical theory suggests:

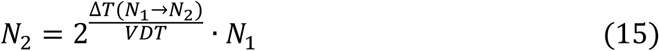

That is,

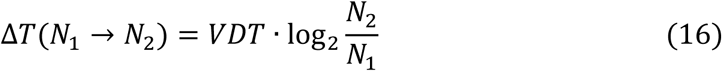

Therefore, from Eq 14:

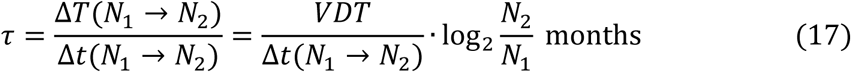

For example, *VDT* is roughly 7 months (211 days) for primary colon cancer^17^. For *N*_1_ of 10^5^ to exponentially grow to *N*_2_ of 10^6^ in primary-tumor simulation in the interval of Δ*t*(10^5^ → 10^6^) simulation time steps,

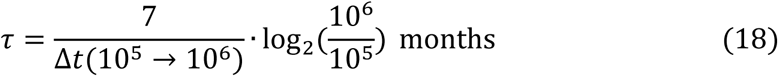

### Drug intervention trial

We implemented the *drug intervention trial* into the program to virtually perform drug intervention. In this trial, the drug efficacy is mimicked by killing a fraction of cells with the malfunction states of user-specified target genes. Suppose that a simulation grows cancer cells at a large volume (*e.g.*, 10^9^ cells) based on best-estimated parameters. Users specify genes to be targeted. When the trial is switched on in the program, the numbers of cells are stochastically sampled for clones with malfunctional target genes in every simulation time step, following a Poisson distribution with the mean derived from a probability specified by the users. Next, the sampled numbers of cells are killed. It is possible to target multiple aberrant genes. The killing probability by virtual drug can be mapped to a real drug dose, by aligning a predicted cell count with the cell count derived from an observed image at one time point after real drug administration (**Supplementary Figure 7b**).

### Modifications to the previous model

We modified the equations (**Supplementary Table 1** and **Supplementary Table 2**), trials (**Supplementary Table 3**), variables (**Supplementary Table 4**), and sampling method of the trials (**Supplementary Figure 9**) proposed in the previous model^14,21^. Each trial related to a cancer hallmark (**Supplementary Table 1**) can now be switched on or off in the program. We have described the modifications in **Supplementary Methods**.

### Generators for parameters and post-processes

We prepared “generators” and post-process programs separately from the main program of tugMedi (**Supplementary Figure 6**). Generators generate random values for parameters used in the main program, based on prior distributions. Such parameters include the simulation time step and waiting division used in the EF algorithm, and the gene type where genes can be oncogenes, suppressors, or dominant-negative genes. Post-process programs calculate, for instance, VAF and TMB from output data from the main program.

A Dirichlet distribution was used as the prior distribution of a generator for the weight parameters of the cell division rate:

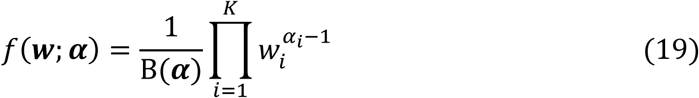

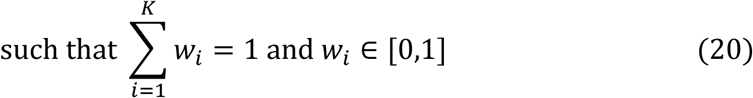

where *f*, ***w***, and α are the probability density function, weight parameters, and Dirichlet α parameters, respectively. B and *K* are the multivariate beta function and number of weight parameters, respectively. Without any prior knowledge, we used the non-informative Dirichlet prior, where all α*_i_* equals one (a uniform distribution over the standard *K*−1 simplex).

### Application to synthetic data

We synthesized VAF and other statistics data based on TCGA 7903 (TCGA ID: TCGA-55-*7903*-01A-11D-2167-08) and AZ (TCGA-*AZ*-6608-01A-11D-1835-10) for synthetic data 1 and 2, respectively. The “true” values of simulation parameters were set arbitrarily, and simulations were conducted using these fixed parameter values. The “true” or “observed” values of statistics, including VAF, were calculated as the averages across 20 simulation replications. From the synthesized VAFs, we estimated parameter values using the same method as used in the real data applications. Then, 2000 simulations were performed using the estimated parameter values for each dataset. Time calibration was performed as in the real data applications. The VDT-based calibration was performed independently for synthesized and simulation data in the synthetic data 2 case.

### Application to TCGA data

From TCGA, we arbitrarily selected patients with quality-passed non-silent SNVs/indels in *APC*, *KRAS*, and *TP53* for colorectal cancer (version: mc3.v0.2.8.PUBLIC.maf) and quality-passed non-silent SNVs/indels in *BRAF* or *EGFR* for lung cancer (version: gdc-1.0.0). We used observed VAFs only in chromosome I as those in other regions for fast computation. We set the cutoff for VAFs in other regions to a minimum of 0.1, based on a conservative limit of detection, and to a maximum corresponding to the largest VAF among driver genes. TMB was calculated based on the selected VAFs. Only for TCGA 8506, we randomly down-sampled poms to 1/20 of the original number because there were too many poms to compute. In the program, we switched only on the cell division trial. CNAs were calculated only for regions of interest. Other simulation settings are provided in settings files in **Data availability** and explained in the vignette of the program in **Code availability**. Parameter estimation is explained in **Supplementary Methods** Ensemble plots were drawn from 2000 simulation replications.

## ABBREVIATIONS

CGP: comprehensive genomic profiling
NGS: next-generation sequencing
CNA: copy number alteration
ITH: intra-tumor heterogeneity
VAF: variant allele frequency
LOH: loss-of-heterogeneity
ABC: approximate Bayesian computation
BO: Bayesian optimization
pom: point mutation
TMB: tumor mutation burden

## Data availability

Simulation settings and results can be accessed at https://figshare.com/s/ce76be8678d2c4d10d62.

## Code availability

tugMedi, version 1.0.32.2 upon submission, can be accessed at https://github.com/tugHall/tugMedi_open. The version used in this study can be accessed in each dataset at the URL provided in **Data availability**.

## Acknowledgments

We thank Munmee Dutta, Towa Fujii, Ritsuko Onuki, and Jo Nishino for their technical assistance and Atsushi Niida for insightful suggestions.

## Author contributions

M.K. designed the study and developed the model. I.N. and M.K. developed the algorithms. I.N., E.F., and M.K. wrote the program codes. M.N. and M.K. prepared the data. E.F. and M.K. performed the analyses. I.N., S.Y., and T.S. reviewed the manuscript. M.K. wrote the manuscript.

## Funding

This work was supported by AMED (22704162 and JP25ym0126804), MEXT (20K12071 and 24K03040), and CREST-JST (JPMJCR 1412). The funders had no role in study design, data collection and analysis, decision to publish, or preparation of the manuscript.

## Competing interests

The authors declare no competing interests.

## Notes

### Competing Interest Statement

The authors have declared no competing interest.

