## Supplementary Information for "tugMedi: simulator of cancer-cell evolution for personalized medicine based on the genomic data of patients"

### SUPPELMENTARY INFORMTATION

#### SUPPLEMENTARY METHODS

##### Equations of hallmark interferences and compaction factors

We arranged and re-formalized the equations of hallmark interferences specified in a previous study[14] in **Supplementary Table 1**. As in a previous study[14], the hallmark variable $H_x$ for hallmark *x* is:

$$H_x = \sum_{i=1}^{n_x} w_{x,i} \cdot g_{x,i} \tag{21}$$

$$\text{such that} \sum_{i=1}^{n_x} w_{x,i} = 1, 0 \leq w_{x,i} \leq 1 \tag{22}$$

$$g_{x,i} = \begin{cases} 1, & \text{when the gene is dysfunctional} \\ 0, & \text{otherwise} \end{cases} \tag{23}$$

where $n_x$ represents the number of genes related to hallmark *x*. $w_{x,i}$ and $g_{x,i}$ are the weight parameter and indicator function indicating the dysfunctional state of gene *i* for hallmark *x*, respectively. $x \in \{\text{d, a, im, i, b}\}$, where d, a, im, i, and b are the hallmark of oncogene/suppressor (summary of self-sufficiency in growth signals and insensitivity to anti-growth signals[42]), apoptosis, invasion/metastasis, immortality, and angiogenesis, respectively. Hallmark variables increase or decrease the differences in trial probabilities from the state of normal cells (**Supplementary Table 1**). **Supplementary Table 2** provides details on some variables in **Supplementary Table 1**. In the simulator, trials related to the hallmarks can be switched on or off individually.

We introduced compaction factor $c_x$ (ranging from 0 to 1) into the equations in **Supplementary Table 1**, because the extent of the interference effect of a hallmark may vary even if all genes related to the hallmark malfunction. Previously[14], all $c_x$ were implicitly assumed at 1. $c_x$ values can be estimated via ABC.

**Modifications to previous trials**

According to the re-formalization in **Supplementary Table 1**, we arranged the trials in **Supplementary Table 3**. We describe below notable points on some trials.

*For the trial of cell division*. The number of cells appearing in the friction term of the logistic growth mode is that of primary-tumor cells (**Supplementary Table 2**). The division rate is dragged through the friction term according to the number of primary-tumor cells (cells with driver mutations) only, excluding normal cells (cells without driver mutations). This friction term is applied to the division rate of both primary-tumor cells and normal cells. In other words, the division rate of both primary-tumor cells and normal cells is dragged only by the number of primary-tumor cells and not influenced by the number of normal cells, because it is reasonably assumed that only primary-tumor cells consume additional resources.

*For the trial of invasion/metastasis transformation*. We changed to this trial based on the “proportional” mode, where the success probability of the invasion/metastasis transformation is proportional to the probability defined for this trial (**Supplementary Table 3**).

*For the trial of replication (Hayflick) limit*. 1) We changed this trial from per time to per division (**Supplementary Table 4**). Immediately after a cell is determined to divide in the cell division trial, the cell division may be canceled depending on the outcome from the replication limit trial. Note that this trial is also applied to normal cells in our simulator, where we postulate that normal cells, such as progenitor cells differentiated from somatic stem cells, cannot indefinitely divide. 2) Performing this trial requires cell division counts per cell in a clone. We modeled their distribution with $Bin(n =$

$N_{\mathrm{divs}}, p = 1/N_{\mathrm{cells}})$, where $Bin(n, p)$ represents the binomial distribution with $n$ Bernoulli trials and the probability of $p$. $N_{\mathrm{divs}}$ and $N_{\mathrm{cells}}$ are the number of cell divisions and the number of cells in a clone at a simulation time step, respectively. We approximated this binomial distribution to a normal distribution, $N\left(\mu = \frac{N_{\mathrm{divs}}}{N_{\mathrm{cells}}}, \sigma^2 = \frac{N_{\mathrm{divs}}}{N_{\mathrm{cells}}} \cdot (1 - \frac{1}{N_{\mathrm{cells}}})\right)$, where $N(\mu, \sigma^2)$ is the normal distribution with mean $\mu$ and variance $\sigma^2$. Taking advantage of the reproductive property, the program computes the distribution of cell division counts per cell in a clone over simulation time steps. 3) We changed the maximum number up to which cells can divide in our model as the maximum number allowed by the Hayflick limit *minus* the number of divisions cells have undergone.

**The Poisson sampling algorithm**

We re-formalized the state-transition process of our model based on a Poisson process, unlike a Bernoulli process previously[14]. For re-formalization, we linked Poisson mean parameters to per-time trial probabilities as follows:

$$\lambda = p' \cdot \kappa \tag{24}$$

where $\lambda$ and $p'$ represent a Poisson parameter and a per-time trial probability, respectively. $\kappa$ is the time-scaling parameter to convert a per-time probability into a

Poisson parameter. $\lambda$ and $p'$ are defined for a single cell. The time unit of $p'$ in simulation is the time interval in which $p'$ events, typically represented with $p'$ cell divisions, occur on average. $\kappa$ scales the time unit as in the Poisson process.

In the computation of our model (**Supplementary Figure 9**), the program randomly draws the number of events occurring during a simulation time step for clone *i* and trial *x* from the Poisson distribution with the mean:

$$n_i \cdot \lambda_{i,x} \tag{25}$$

where $n_i$ is the number of cells in clone *i*, and $\lambda_{i,x}$ is the same as Eq. 24, *i.e.*, the Poisson parameter for a single cell. $x \in$ {d, k, a, im}, where d, k, a, and im are the trials of cell division, constant cell death, apoptosis, and invasion/metastasis transformation, respectively.

According to the drawn number, the program executes the trial events to change the state of a clone (**Supplementary Figure 9**). For example, if 3 is sampled from the Poisson distribution with $n_i\lambda_{i,\mathrm{k}}$ in clone *i* for constant cell death trial k, 3 cell death events occur, leading to a change in clone size $n_i$ to $n_i$ - 3 for this trial. Depending on a trial, other non-per-time trials and processes are successively performed, such as the gene mutation trial (*i.e.*, mutations may occur) (**Supplementary Figure 9**).

If cell divisions occur, new cells are generated; however, we do not perform the operation above for new cells until the next simulation time step. We prohibit biologically contradictory serial events for a single cell: for example, 3 death events may occur during a time step on a single cell, but we only allow one death event for the cell. Finally, changing the state of all clones from $S_{\mathrm{b}}$ (before) to $S_{\mathrm{a}}$ (after), we carry forward the simulation time by one (**Supplementary Figure 9**). We repeat this state-change process during a simulation.

**Equilibrium state of normal cells**

We assume equilibrium between birth and death for normal cells, such that the number of normal cells is constant on average. If users do not specify a specific value in the simulator, constant death rate parameter $k_{\mathrm{N}}$ is set to result in this equilibrium. Such $k_{\mathrm{N}}$ is derived as follows: consider only the constant death, apoptosis, and cell division trials for normal cells, because the invasion/metastasis trial does not apply to normal cells and the replication limit trial only works after dozens of cell divisions. The change, $\Delta N$, in normal cell numbers from the state before to after a simulation step is on average:

$$\Delta N = \sum_{i=1}^{N_\mathrm{C}} n_i \left(\lambda_{i,\mathrm{d}} - \lambda_{i,\mathrm{a}} - \lambda_{i,\mathrm{k}}\right) \tag{26}$$

$$= \kappa \sum_{i=1}^{N_\mathrm{C}} n_i \left(d_{\mathrm{N},i} - a_{\mathrm{N},i} - k_\mathrm{N}\right) \tag{27}$$

where $N_\mathrm{C}$ is the number of clones comprising normal cells and $n_i$ is the number of normal cells in clone *i*. $\lambda_{i,\mathrm{d}}$, $\lambda_{i,\mathrm{a}}$, and $\lambda_{i,\mathrm{k}}$ are the Poisson parameters for the numbers of events of the three trials (d, cell division; a, apoptosis; k, constant death trials) for a single normal cell; and $\kappa$, and $d_{\mathrm{N},i}$ and $a_{\mathrm{N},i}$ are the time-scaling parameter to obtain the Poisson parameters, and the original binomial probabilities of the two trials, respectively. Because $d_{\mathrm{N},i}$ and $a_{\mathrm{N},i}$ are the same across normal clones:

$$\Delta N = \kappa(d_\mathrm{N} - a_\mathrm{N} - k_\mathrm{N}) \sum_{i=1}^{N_\mathrm{C}} n_i \tag{28}$$

In the equilibrium state, *i.e.*, $\Delta N = 0$:

$$k_\mathrm{N} = d_\mathrm{N} - a_\mathrm{N} \tag{29}$$

$k_\mathrm{N}$ determined using this equation balances the birth and death of normal cells. $a_\mathrm{N}$ is a small value in the default setting (~$6.7\times10^{-3}$, with $s_\mathrm{N} = 10$ and $x = 0$). When $a_\mathrm{N}$ is regarded as zero,

$$k_\mathrm{N} = d_\mathrm{N} \tag{30}$$

##### $\boldsymbol{\rho}_{\mathrm{N}}$ estimation using the VAF equation

$\rho_{\mathrm{N}}$, the admixture rate of intact normal cells in Eq. 5, is an external parameter for the simulator. It is estimated from values given by users or values sampled from a prior distribution, by evaluating the fit of VAFs between simulated and observed values. Given a value $\rho'_{\mathrm{N}}$, simulated VAFs can be calculated from simulated $n^i_{\mathrm{A},s}$ and $n^i_{\mathrm{B},s}$, and $\tau_s$ using Eq. 5. However, $\tau_s$ needs to be scaled from $\lambda_s$, which represents the fraction of a subpopulation comprising speckled normal and tumor cells in a simulation such that

$$\sum_{s=1}^{\#sp} \lambda_s = 1 \tag{31}$$

(**Supplementary Figure 5b**), because our simulator does not simulate normal cell contamination. Under unbiased sampling, we can assume:

$$\tau_1 : \tau_2 : \cdots = \lambda_1 : \lambda_2 : \cdots \tag{32}$$

Thus,

$$\tau_s = k\lambda_s \tag{33}$$

where *k* is a scaling parameter. From Eq. 4,

$$\sum_{s=1}^{\#sp} \tau_s + \rho'_{\mathrm{N}} = k \sum_{s=1}^{\#sp} \lambda_s + \rho'_{\mathrm{N}} = k + \rho'_{\mathrm{N}} = 1 \tag{34}$$

Hence,

$$k = 1 - \rho'_{\mathrm{N}} \tag{35}$$

That is,

$$\tau_s = (1 - \rho'_{\mathrm{N}})\lambda_s \tag{36}$$

Therefore, $\tau_s$ can be calculated from $\lambda_s$ of a simulation (**Supplementary Figure 5b**), yielding simulated VAFs under $\rho'_{\mathrm{N}}$. Simulated VAFs are then used to evaluate the fit with observed VAFs.

Alternatively, a tumor purity pathologically measured may be given, say, $\rho'_{\mathrm{T}}$. From Eq. 2:

$$\rho'_{\mathrm{T}} = \sum_{s \in \mathrm{T}} \tau_s = k \sum_{s \in \mathrm{T}} \lambda_s \tag{37}$$

Thus, from

$$k = \rho'_{\mathrm{T}} \Big/ \sum_{s \in \mathrm{T}} \lambda_s \tag{38}$$

$\tau_s$ can be calculated from $\lambda_s$ of a simulation as:

$$\tau_s = k\lambda_s = \left(\rho'_{\mathrm{T}} \Big/ \sum_{s \in \mathrm{T}} \lambda_s\right)\lambda_s \tag{39}$$

In this study, we estimated $\rho_{\mathrm{N}}$ with Eq. 36. Note that the subpopulations of speckled normal cells are included in VAF calculation for primary tumors but excluded from metastatic tumors, because it is unlikely for speckled normal cells to metastasize.

**Parameter estimation by ABC**

For ABC, we used the abc package in R, as used previously[14,33]. We used VAFs as summary statistics in ABC. Since the observed VAFs were from primary tumors, we compared observed VAFs with VAFs calculated from primary tumor cells in simulations in this study. Because roughly $10^8$–$10^9$ cancer cells are at the clinically detection level of cancer[41], we used VAFs at the simulation time step when the number of cancer cells reached $10^9$ for ABC.

For VAF comparison, in each gene, we first sorted VAFs in descending order for each of the observed and simulated VAFs. Next, we made the pairs of observed and simulated VAF values from highest to lowest values. If a VAF was vacant in a pair, we supplemented a value of 0. For example, if there were 10 simulated VAFs and 7 observed VAFs, we set 8–10th observed VAFs to 0. We changed VAF values less than 0.1 to 0, using 0.1 as the conservative limit of detection in NGS. We discarded paired VAFs of which both values were 0. We treated other regions collectively as one gene because these are indistinguishable.

ABC selects input parameter values that yield the simulated values of summary statistics close to observed values (**Supplementary Figure 6**). These input parameter values can be directly resampled or summarized to update hyper-parameters for the next-round simulations (**Supplementary Figure 6**). Using both strategies, we iterated parameter updating several times until a good fit was obtained.

Specifically, we estimated the mutation rate $m_0$, the base rate of cell divisions $d_N$ ($k_N$ was set to lead to the equilibrium state), the weight parameters of the cell division rate, and the simulation time steps for mutation occurrences, employing the following priors at the first iteration.

- $m_0$: discrete uniform distribution over $\{10^{-7}, 10^{-8}, 10^{-9}, 10^{-10}\}$
- $d_N$: discrete uniform distribution over $\{10^{-3}, 10^{-4}, 10^{-5}, 10^{-6}, 10^{-7}\}$
- Weight parameters: non-informative Dirichlet distribution
- Time steps
  - First-hit mutation: 1
  - Second-hit mutation: uniform distribution over 2 – 150 in integers
  - Third-hit mutation: uniform distribution over 3 – 150 in integers

  where 150 corresponds to the time step when the number of cells reached $10^9$ in most cases.

Simulations were performed 2000 times, and simulation replications were selected based on the rejection algorithm with a 5% tolerance rate. The selected replications were then randomly resampled for the second iteration.

At the second iteration, simulations were performed 2000 times and a data frame comprising the ABC distance and the parameters was obtained. Bayesian optimization (BO) was applied to the data frame, as described below, to obtain the point estimates of the parameters. When a good VAF fit was not achieved, we manually fine-tuned the estimates. For example, when $m_0$ was estimated as $10^{-5.2}$, the prior was updated to a narrowed uniform distribution of $\{10^{-5}, 10^{-6}\}$ for the third iteration. Similarly, the mutation timing was adjusted to a narrowed uniform distribution around the point estimate (*e.g.*, ±20) for the third iteration. Then 2000 simulations were performed in the same way. This fine-tuning was repeated until a good fit was achieved. The program codes and settings are provided in **Code availability** and **Data availability**, respectively.

**Parameter estimation by BO**

To obtain a point estimate from a joint posterior distribution implied by points selected in ABC, we applied BO to the data frame described above in ABC, where the ABC distance was treated as the response variable to be minimized. Specifically, we first obtained a score function representing the ABC distance by performing Gaussian process regression using the km() function in the DiceKriging library of R. $m_0$ and $d_N$ were log-transformed. We then searched for the optimal point that minimizes the score function, using the BayesianOptimization() function in the rBayesianOptimization library of R. The program codes and settings are provided in **Code availability**.

#### Evaluation of data fit

We evaluated data fitting of a summary statistic by the mean error (*ME*):

$$ME = s^{\mathrm{obs}} - \frac{1}{n}\sum_{r=1}^{n} s_r^{\mathrm{sim}} \tag{40}$$

where *s* is a summary statistic, and “obs” and “sim” represent observed and simulated values, respectively. *n* is the number of simulation replications. As the summary statistics, we used observed and simulated VAFs that were paired as described above in the subsection on ABC. We used the limit of detection described above as the cut-off for the observed and simulated VAFs.

**Implementation notes on CNA processes**

Herein, we explain how we handled deletions and duplications along a chromosome computationally. For convenience, we distinguished the *reference* genome position coordinate system from the genome position coordinate system of an *individual* patient, termed reference and physical positions, respectively. The reference genome does not shrink or extend, but the patients' physical genomes do.

For example, in the *event* step at time $t$ in **Supplementary Figure 3a**, a deletion occurs with the breakpoints of the physical positions of 4 and 6 (length 3) on a parental chromosome. In the *effect* step, this deletion removes the physical positions of 4, 5, and 6. Lastly, in the *renumber* step, physical positions are simply renumbered from left to right. Consequently, although the reference positions are invariant, the physical positions are changed and the total physical length shrinks.

Next at time $t + 1$, in the event step, another deletion occurs with the breakpoint positions of 3 and 5 (length 3) in the physical position coordinate system. In the effect step, this deletion removes the physical positions of 3, 4, and 5. In the renumber step, the physical positions are renumbered. Thus, again the reference positions are not changed but the physical positions are changed and the physical length shrinks further.

To calculate the copy number, we introduced a convenient notation, $B^{t}_{r(p)}$, where B represents variant allele B, and *t*, *r*, and *p* represent time *t*, the reference, and physical positions where the B is located, respectively. Let $B^{t}_{5(5)}$ located at the reference and physical positions of 5 and 5 at time *t* in **Supplementary Figure 3a**. In the renumber step, it is renumbered to $B^{t}_{5(\text{-})}$, where "-" represents a deletion. Meanwhile, $B^{t}_{9(9)}$ changes to $B^{t}_{9(6)}$. In the renumber step at time *t* + 1, the former variant is changed only for the time, $B^{t+1}_{5(\text{-})}$, whereas the latter variant is changed for the physical position and time, $B^{t+1}_{9(3)}$. The reference positions of both variants do not change.

As shown in **Supplementary Figure 3b**, a duplication event occurs with the breakpoints of the physical positions of 4 and 6 (length 3) on a parental chromosome at time *t*. In the effect step, a chromosomal segment between the breakpoints is duplicated in the physical and reference position coordinates as 4-5-6 and 4-5-6. In the renumber step, the numbers of the duplicated and subsequent physical positions are renumbered from left to right. Consequently, the physical positions become unique in ascending order, while the reference positions are repeated as 4-5-6 and 4-5-6. This means, for example, physical positions 6 and 9 originate from reference position 6. The total physical length is extended to 12.

Next at time *t* + 1, in the event step, another duplication event occurs with the breakpoint positions of 9 and 10 (length 2) in the physical position coordinate system. In the effect step, a chromosomal segment between the breakpoints is duplicated in the physical position coordinate as 9-10 and 9-10 and the reference coordinate as 6-7 and 6-7. In the renumber step, the physical positions are renumbered. Thus, again the physical positions are unique in ascending order, while the reference positions are repeated as 6-7 and 6-7. Physical positions 6, 9, 11 originate from reference positions 6. The total physical length is extended to 14.

Variant $B^{t}_{6(6)}$ is duplicated into $B^{t}_{6(6)}$ and $B^{t}_{6(9)}$ in the renumber step at time *t*, as shown in **Supplementary Figure 3b**. Due to the second duplication at time *t* + 1, variant $B^{t}_{6(9)}$ is duplicated into $B^{t+1}_{6(9)}$ and $B^{t+1}_{6(11)}$ in the renumber step. After all, $B^{t}_{6(6)}$ is duplicated into $B^{t+1}_{6(6)}$, $B^{t+1}_{6(9)}$, and $B^{t+1}_{6(11)}$. Meanwhile, variant $B^{t}_{9(9)}$ is not duplicated at the renumber step at time *t* or *t* + 1 but due to the two duplication events, the physical position is changed to $B^{t}_{9(12)}$ at time *t* and $B^{t+1}_{9(14)}$ at time *t* + 1. The reference positions of all variants derived from $\mathrm{B}^{t}_{6(6)}$ and $\mathrm{B}^{t}_{9(9)}$ do not change at *t* + 1.

The copy number of a variant is easily converted from notation $\mathrm{B}^{t}_{r(p)}$. Let us define $cn(\mathrm{B}^{t}_{r(p)})$ as the copy number of $\mathrm{B}^{t}_{r(p)}$. For example, $cn\left(\mathrm{B}^{t}_{6(6)}\right) = 1$ in the event step at *t,* as shown in **Supplementary Figure 3c**. When the physical position is "-" the copy number of a variant is 0. For example, in the renumber step at *t*, $cn\left(\mathrm{B}^{t}_{6(\text{-})}\right) = 0$, because the variant is removed from the chromosome by the deletion event. Meanwhile, $cn\left(\mathrm{B}^{t+1}_{8(5)}\right) = 1$ at the event step at *t* + 1. Then, the physical position is removed in the effect step, and $cn\left(\mathrm{B}^{t+1}_{8(\text{-})}\right) = 0$.

In the event step at *t*, as shown in **Supplementary Figure 3d**, $cn\left(\mathrm{B}^{t}_{6(6)}\right) = 1$. This variant is duplicated, and the duplicates with the same reference position are summarized as $\mathrm{B}^{t}_{6(6,9)}$ in the renumber step. Since this notation has two physical positions, $cn\left(\mathrm{B}^{t}_{6(6,9)}\right) = 2$. At *t* + 1, $\mathrm{B}^{t+1}_{6(9)}$ is duplicated to $\mathrm{B}^{t+1}_{6(9)}$ and $\mathrm{B}^{t+1}_{6(11)}$. All the duplicated variants with the same reference position are summarized as $\mathrm{B}^{t+1}_{6(6,9,11)}$. Thus, $cn\left(\mathrm{B}^{t+1}_{6(6,9,11)}\right) = 3$. The same logic applies to the original allele noted as $\mathrm{A}^{t}_{r(p)}$, where A represents the original allele. Suppose $cn\left(\mathrm{A}^{t+1}_{6(2,4)}\right) = 2$. VAF at reference position 6 at *t* + 1 is calculated from these copy numbers (3 for B and 2 for A).

#### SUPPLEMENTARY TABLES

**Supplementary Table 1.** Equations for hallmark interferences, and compaction factors

| Trial[14] | Hallmark in tugHall[14] | Re-formalized equation | |
|---|---|---|---|
| Cell division | Oncogene/suppressor | $d' - d_{\mathrm{N}} = +c_{\mathrm{d}} H_{\mathrm{d}} \Gamma$ | (41) |
| Apoptosis | Apoptosis | $a' - a_{\mathrm{N}} = -c_{\mathrm{a}} H_{\mathrm{a}}$ | (42) |
| Invasion/metastasis transformation | Invasion/metastasis | $im' - im_{\mathrm{N}} = +c_{\mathrm{im}} H_{\mathrm{im}}$ | (43) |
| Replication (Hayflick) limit | Immortality | $i' - i_{\mathrm{N}} = -c_{\mathrm{i}} H_{\mathrm{i}}$ | (44) |
| Cell division via carrying capacity, *K* | Angiogenesis | $\log_{10} K' - \log_{10} K_{\mathrm{N}} = +\frac{1}{c_{\mathrm{b}}} H_{\mathrm{b}} = +F_{\mathrm{b}} H_{\mathrm{b}}$ | (45) |

We re-formalized equations describing the relationships between trial probabilities (or a related variable: carrying capacity) and hallmark variables. Hallmark variables influence the differences in trial probabilities from the state of normal cells subscripted by N. See **Supplementary Table 4** for variable notations. Variables with prime (‘) are probabilities actually used for trials. Compaction factor $c_x$ ($x \in$ {d, a, im, i, b}), ranging from 0 to 1, is introduced in this study. *K* and $F_{\mathrm{b}}$ in Eq. 45 denote the carrying capacity of the logistic growth in cell division and the expansion factor in log for carrying capacity, respectively. This equation is equivalent to $K'/K_{\mathrm{N}} = 10^{F_{\mathrm{b}} H_{\mathrm{b}}}$. See **Supplementary Table 2** for detailed notes on some variables.

**Supplementary Table 2.** Detailed notes on variables in the hallmark equations

| Hallmark equation | Variable to note | Note |
|---|---|---|
| $d' - \boldsymbol{d}_{\mathrm{N}} = +c_{\mathrm{d}} H_{\mathrm{d}} \boldsymbol{\Gamma}$ | $d_{\mathrm{N}}$ | Constant parameter, given by users |
| | $\Gamma$ | $\Gamma = \begin{cases} \Gamma_{\mathrm{l}} = 1 - N_{\mathrm{P}}/K', & \text{for logistic growth mode} \\ \Gamma_{\mathrm{e}} = 1, & \text{for exponential growth mode} \end{cases}$ (46) |
| $a' - \boldsymbol{a}_{\mathrm{N}} = -c_{\mathrm{a}} H_{\mathrm{a}}$ | $a_{\mathrm{N}}$ | $a_{\mathrm{N}} = \sigma\left(s_{\mathrm{N}} \times (x - 0.5)\right)$ (47)<br>where $\sigma$ is a sigmoid function defined by parameter $s_{\mathrm{N}}$,<br>$x$ is TMB, i.e., the number of passenger poms per Mb |
| $im' - \boldsymbol{im}_{\mathrm{N}} = +c_{\mathrm{im}} H_{\mathrm{im}}$ | $im_{\mathrm{N}}$ | Constant, practically 0 |
| $i' - \boldsymbol{i}_{\mathrm{N}} = -c_{\mathrm{i}} H_{\mathrm{i}}$ | $i_{\mathrm{N}}$ | Constant, practically 1 |
| $\log_{10} K' - \log_{10} \boldsymbol{K}_{\mathrm{N}} = +\frac{1}{c_{\mathrm{b}}} H_{\mathrm{b}}$ | $K_{\mathrm{N}}$ | Constant parameter, given by users |

$N_{\mathrm{P}}$ in Eq. 46 is the number of primary-tumor cells. Saturated number of cells in the logistic growth is different from *K'* when cells are influenced by trials other than the cell division trial.

**Supplementary Table 3.** The trials

| Trial | Condition | Probability | Event |
|---|---|---|---|
| **Constant death** | Every time step | ● $k' = k_{\mathrm{N}}$ | Death |
| | | ● $1 - k'$ | Nothing |
| Apoptosis | Every time step | ● $a'$ | Death |
| | | ● $1 - a'$ | Nothing |
| Invasion/ metastasis transformation | $im' \neq 0$ | **Proportional mode** | |
| | | ● $\boldsymbol{im' \cdot Z}$ | **Transform into exponential growth** |
| | | ● $\boldsymbol{im' \cdot (1 - Z)}$ | **Death** |
| | | ● $\boldsymbol{1 - im'}$ | **Nothing** |
| Replication limit | $ct > ct_{\mathrm{max}}$ | ● $i'$ | Stop division process |
| | | ● $1 - i'$ | Start division trial |
| Cell division | Every time step | ● $d'$ | Division |
| | | ● $1 - d'$ | Nothing |
| Gene mutation | Cell division happens | ● Poisson distribution with the mean of $m_{0,\mathrm{pom}} \times$ *length_exon* | Point mutation |
| | | ● **Poisson distribution with the mean of $\boldsymbol{m_{0,\mathrm{del}}} \times$ *length_exon_intron*** | **Deletion breakpoint** |

| | | | |
|---|---|---|---|
| | | • **Exponential distribution with the mean of** $\boldsymbol{l_{0,\mathrm{del}}}$ | **Deletion length** |
| | | • **Poisson distribution with the mean of** $\boldsymbol{m_{0,\mathrm{dup}}} \times$ ***length_exon_intron*** | **Duplication breakpoint** |
| | | • **Exponential distribution with the mean of** $\boldsymbol{l_{0,\mathrm{dup}}}$ | **Duplication length** |
| Gene malfunction | Gene mutation happens | • $u'_{\mathrm{pom}} = \begin{cases} u_{0,\mathrm{pom,o}}, \text{ for oncogene} \\ u_{0,\mathrm{pom,s}}, \text{ for suppressor} \end{cases}$ | Malfunction by point mutation |
| | | • $1 - u'_{\mathrm{pom}}$ | Nothing |
| | | • $\boldsymbol{u'_{\mathrm{del}} = \begin{cases} u_{0,\mathrm{del,o}}, \text{ for oncogene} \\ u_{0,\mathrm{del,s}}, \text{ for suppressor} \end{cases}}$ | **Malfunction by deletion** |
| | | • $\boldsymbol{1 - u'_{\mathrm{del}}}$ | **Nothing** |
| | | • $\boldsymbol{u'_{\mathrm{dup}} = \begin{cases} u_{0,\mathrm{dup,o}}, \text{ for oncogene} \\ u_{0,\mathrm{dup,s}}, \text{ for suppressor} \end{cases}}$ | **Malfunction by duplication** |
| | | • $\boldsymbol{1 - u'_{\mathrm{dup}}}$ | **Nothing** |
| **Drug intervention** | **Every time step (during user-specified intervals)** | **For user-specified target genes** | |
| | | • $\boldsymbol{DI_{\mathrm{k}}}$ | **Killing cells with malfunctional genes** |
| | | • **1-**$\boldsymbol{DI_{\mathrm{k}}}$ | **Nothing** |

We modified Supplementary Table 2 from [14]. Notable changes are indicated in bold. The "condition" column represents the condition to which the trial is applied. See **Supplementary Table 4** for variable notations. The trials of gene mutation and gene malfunction are implemented using the EF algorithm.

**Supplementary Table 4.** The variables

| Variable type | Notation | Description | Per | Interfered by hallmarks | Time change |
|---|---|---|---|---|---|
| Cell | $ct$ | Cell division counter | - | - | Dynamic |
| | $ct_{max}$ | **Residual number to the maximum cell division number of replication limit** | - | - | Static |
| | $k$ | Rate of **constant** cell death | Time | - | Static |
| | $d$ | Rate of cell division | Time | Yes | Dynamic |
| | $im$ | Rate of invasion/metastasis transformation | Time | Yes | Dynamic |
| | $a$ | Rate of cell death by apoptosis | Time | Yes | Dynamic |
| | $i$ | Probability of cell division stop by replication limit | **Division** | Yes | Dynamic |
| | $m_{pom}$ | Point mutation rate per bp | Division | - | Static |
| | $\boldsymbol{m_{del}}$ | **Deletion (breakpoint) rate per bp** | **Division** | **-** | **Static** |
| | $\boldsymbol{m_{dup}}$ | **Duplication (breakpoint) rate per bp** | **Division** | **-** | **Static** |
| | $\boldsymbol{l_{del}}$ | **Mean deletion length in bp** | **Del** | **-** | **Static** |
| | $\boldsymbol{l_{dup}}$ | **Mean duplication length in bp** | **Dup** | **-** | **Static** |
| | $u_{pom,o}$, $u_{pom,s}$ | Probability of malfunction of an oncogene/suppressor by point mutation | Pom | - | Static |
| | $\boldsymbol{u_{del,o}}$, $\boldsymbol{u_{del,s}}$ | **Probability of malfunction of an oncogene/suppressor by deletion** | **Del** | **-** | **Static** |

| | | | | | |
|---|---|---|---|---|---|
| | **$u_{dup,o}$, $u_{dup,s}$** | **Probability of malfunction of an oncogene/suppressor by duplication** | **Dup** | **-** | **Static** |
| External | **$N_N$, $N_{SN}$** | **Number of intact/speckled normal cells (logistic growth)** | **-** | **-** | **Dynamic** |
| | $N_P$ | Number of primary-tumor cells (logistic growth) | - | - | Dynamic |
| | $N_M$ | Number of metastatic-tumor cells (exponential growth) | - | - | Dynamic |
| | $K$ | Carrying capacity in logistic growth | - | - | Static |
| | ($E$ | = 1/$K$, environmental resource limitation) | | | |
| | $F$ | Expansion order to $K$ by angiogenesis | - | Yes | Static |
| | **$Z$** | **Success rate in invasion/metastasis transformation for cells on the trial** | **Time** | **-** | **Static** |
| | **$\kappa$** | **Time-scaling parameter to convert trial probabilities into Poisson lambdas, $\lambda = \kappa p$** | **-** | **-** | **Static** |
| | $T$ | Time counter | - | - | - |
| **Intervention** | **$DI_k$** | **Probability of killing cells with gene aberrations in drug intervention.** | **Time** | **-** | **Static** |

We modified Supplementary Table 1 from [14]. Notable changes are indicated in bold. Cell variables in the "variable type" column are those attributed to each cell and inherited by the daughter cell in cell division. External variables are those that represent the outside environments of cells. The intervention variable represents a variable related to drug intervention. The "Per" column represents units by which the rates or probabilities are defined. Dynamic and static variables in the “time change” column denote whether values change or not with time, respectively. The initial values of some variables need to be specified to start a simulation. See the vignette of the software for specific values and the corresponding reasons.

#### SUPPLEMENTARY FIGURE LEGENDS

**Supplementary Figure 1.** Illustration of the EF algorithm.

**(a)** For regions of interest. 1) Before starting a simulation, a user specifies M1, M2, and M3, which are possible driver mutations (*e.g.*, in *APC*, *KRAS*, and *TP53*, respectively). M2 and M3 are conditioned to occur in cells with M1, and M1 and M2, respectively. 2) Simulation time steps are randomly sampled from prior distributions a user specifies. 3) A simulation starts. Along the simulation run, the mutations are enforced to occur in cells randomly selected at the sampled time steps (letters in red). For M2 and M3, such cells are selected from cells with M1, and M1 and M2, respectively (letters in orange). **(b)** For other regions. 1) Before starting a simulation, a user specifies the number of passenger mutations. 2) Either pom, del, or dup is randomly selected according to the relative probabilities calculated from the Poisson $\lambda$ parameters summed across regions. Next, specific regions (typically, genes) are randomly selected according to the sizes of the regions (*i.e.*, according to $l_i$ and $L_i$: the total exon and exon + intron lengths of region *i*, respectively). 3) Waiting divisions are randomly sampled from exponential distributions with the corresponding Poisson parameters. 4) Mutations are inserted into cells at the division timing specified by the sampled waiting divisions (letters in red). **(c)** The position of a generated CNA is uniform-randomly determined to overlap with region *i*.

**Supplementary Figure 2.** CNA processes in simulation

**(a)** Changes in simulation processes related to CNAs from previous studies[14,21]. **(b)** Changes focusing on point mutations. P1 and P2 represent parental 1 and 2, respectively. **(c)** Representative mechanisms of CNA generation. **(d)** Simulation processes of CNAs. **(e)** Model of gene dysfunction.

**Supplementary Figure 3.** Physical and reference positions representing CNAs. See **Supplementary Methods** for details. **(a)** Physical and reference positions influenced by deletion events. **(b)** Physical and reference positions influenced by duplication events. **(c)** Copy numbers of variants influenced by deletion events. **(d)** Copy numbers of variants influenced by duplication events.

**Supplementary Figure 4**. Genetic modes.

**(a)** In the recessive mode, only when a gene in both parental chromosomes malfunctions, the cell division rate (generally, hallmark variables in **Supplementary Methods**) is affected. In the dominant mode, the malfunction state in one parental chromosome is sufficient for the effect. **(b)** Poms and dups in oncogenes follow the dominant mode, but dels do not have any effect. Poms and dels in suppressors follow the recessive mode, but dups do not have any effect. Dominant-negative genes are the same as suppressors, except for poms in dominant-negative genes following the dominant mode. These rules are implemented in the program. *u* represents the malfunction rate, and the subscripts "o" and "s" represent oncogene and suppressor, respectively.

**Supplementary Figure 5.** Three cell types and their implication in the VAF equation. **(a)** Three cell types and relevant notations in the VAF equation. Intact normal cells are cells without any mutations (poms or CNAs). Speckled normal cells are cells with passenger mutations but without driver mutations. Tumor cells are cells with one or more driver mutations. The pom represented as allele B at site *i* is a passenger mutation. **(b)** Relationship between $\tau_s$ and $\lambda_s$ in the estimation of $\rho_{\mathrm{N}}$.

**Supplementary Figure 6. Generators, post-processes, and ABC/BO.**

Generators generate sets of random values from prior distributions with hyper-parameters. Such sets are indicated as Input 1, Input 2, ... These sets are used as input parameter values in the main tugMedi program, which then returns the output of respective sets of output data, indicated as Output 1, Output 2, .... Post-process programs process the output data to output additional data, such as VAFs at $\rho_1$, VAFs at $\rho_2$, ..., where $\rho_i$ represents the normal-cell admixture rate $i$, such as 0%, 10%, 20%, ... ABC selects input sets yielding the simulated values of summary statistics (VAFs) close to observed values. $\rho_i$ is mainly selected via ME, the mean error (**Supplementary Methods**). The selected input values are directly resampled, or used for BO and manual fine-tuning to update hyper-parameters for the next simulation.

**Supplementary Figure 7.** Calibration of time and drug killing rate.

**(a)** Conversion of a simulation time unit into a real time unit. The settings in the figure are a virtual example for illustration. The real tumor diameters and time interval are derived from a patient with lung cancer[16]. We first calculate the volumes of cells from the diameters with the formula for the volume of a spheroid: $\left(\frac{4\pi}{3}\right)\cdot\left(\frac{L}{2}\right)\cdot\left(\frac{S}{2}\right)\cdot\left(\frac{S}{2}\right) \sim \boldsymbol{L}\cdot\boldsymbol{S}\cdot\boldsymbol{S}/\boldsymbol{2}$, where $L$ and $S$ are long and short diameters ($\boldsymbol{L}=\boldsymbol{S}$ in this case), respectively. Then we obtain the number of cells through the conversion rate of $10^6$ cells per 1 $mm^3$ [41]. We find a simulation time interval yielding the same numbers of cells in simulation and scale the interval to the real time interval. **(b)** Mapping a drug killing rate to the real drug dose. Prediction curves are first plotted with multiple possible drug-killing rates (broken lines). The drug dose is then mapped to a drug killing rate that matches the cell count derived from an observed radiological image (red point). The drug dose is mapped to a killing rate of 60% in this figure.

**Supplementary Figure 8.** Comparison between true and estimated parameter values

The *x*-axis indicates the parameters, and the *y*-axis indicates the values, covering a scaled search space. The circle in red (with solid line) and cross in blue (with broken line) represent the true and estimated values, respectively. **(a)** For synthetic data 1. **(b)** For synthetic data 2.

**Supplementary Figure 9.** Illustration of the sampling algorithm of the trials.

The algorithm is based on sampling from Poisson distributions. The number of occurrences is sampled with a Poisson parameter and the state of a clone is changed according to the sampled number of a trial. $S_b$ and $S_a$ represent the states before and after this process, respectively. These trials are performed independently, and users can switch on or off each of them in the program.

### Supplementary Figure 1

(a)

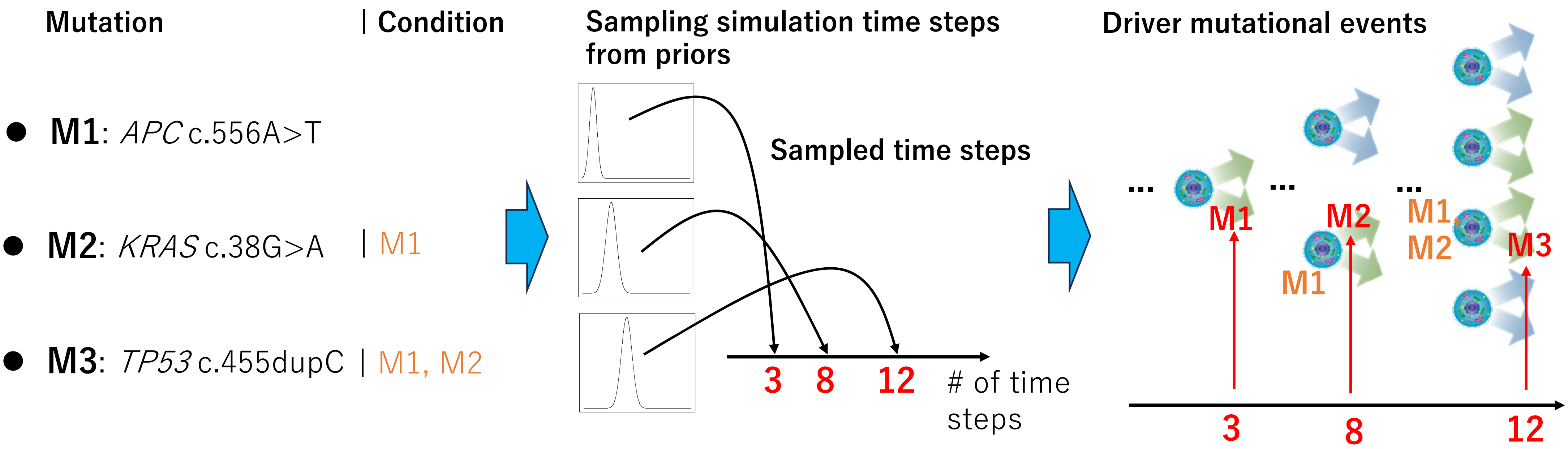

### Supplementary Figure 1 (cont'd)

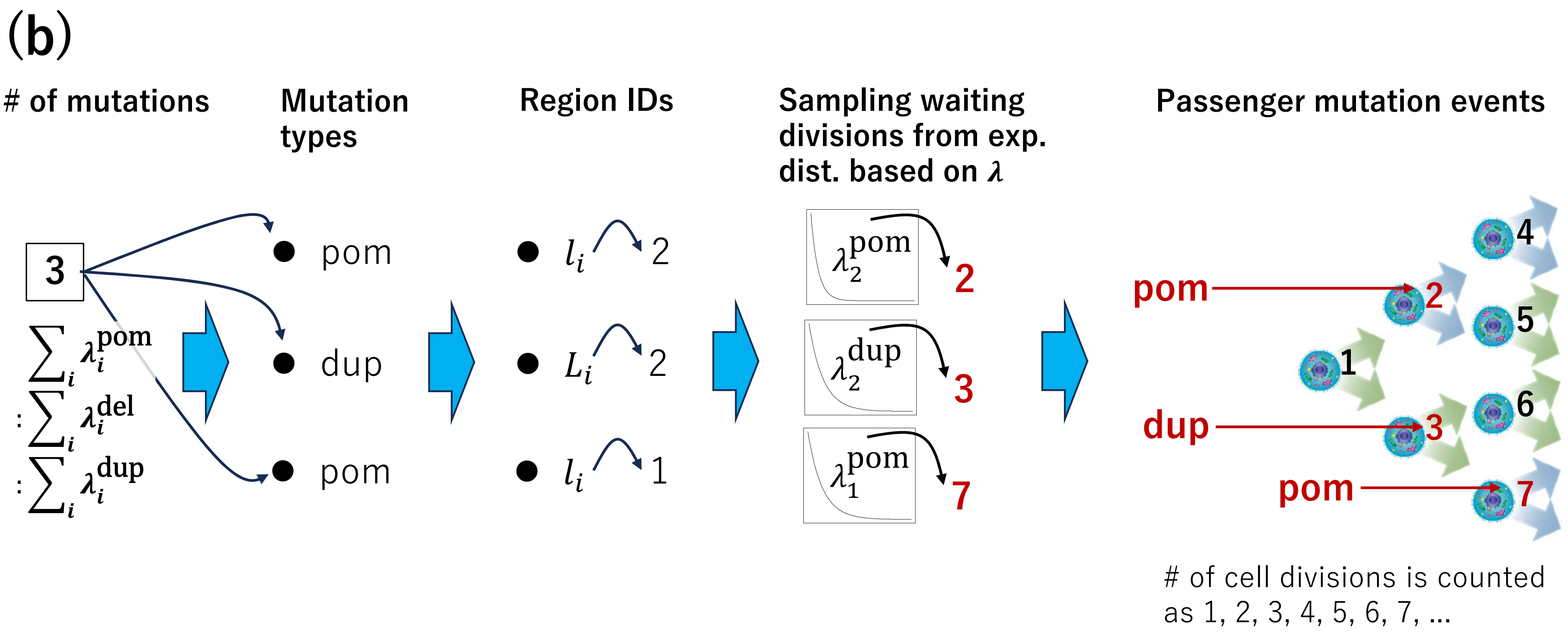

### Supplementary Figure 1 (cont'd)

(c)

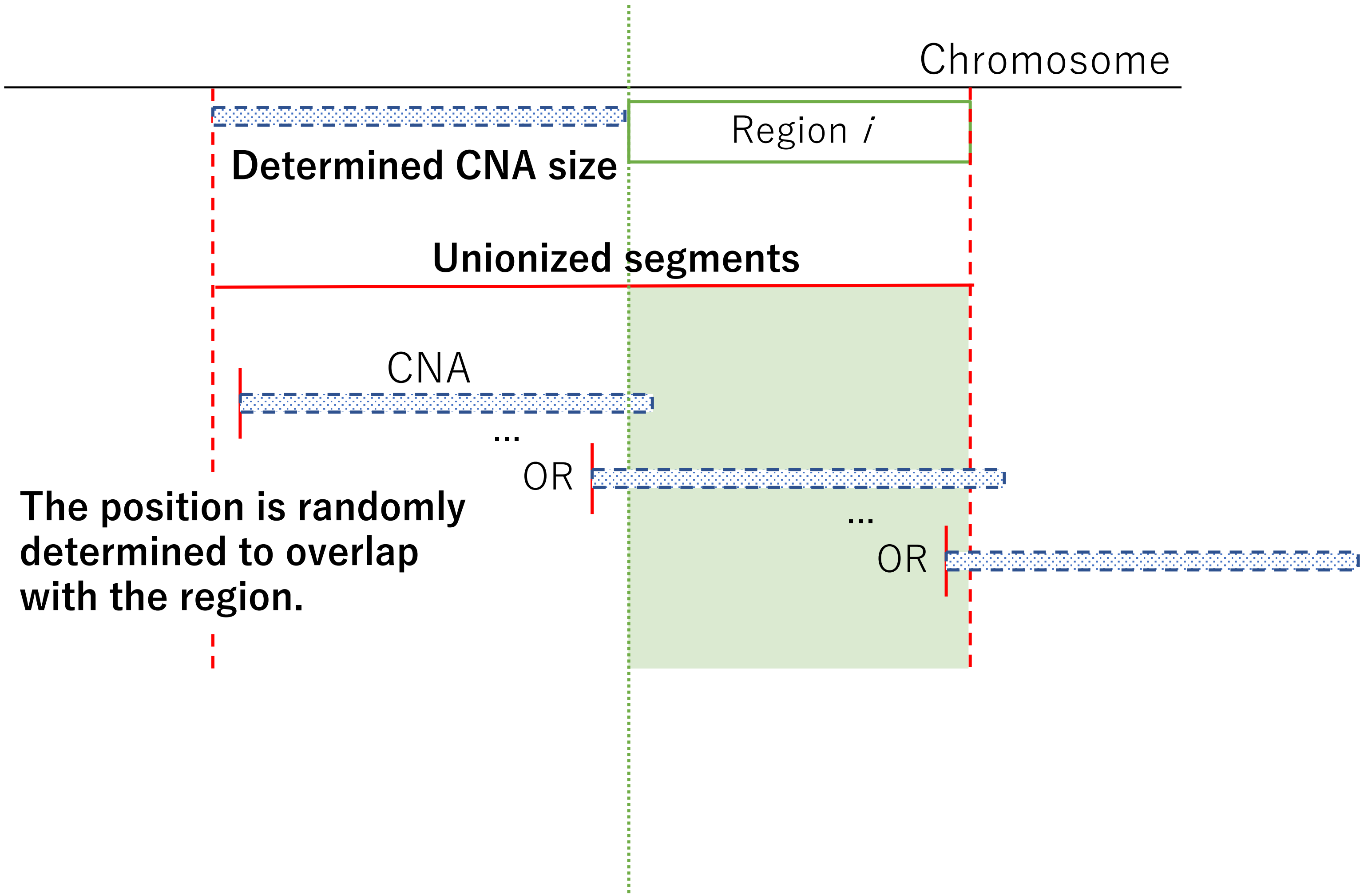

### Supplementary Figure 2

(a)

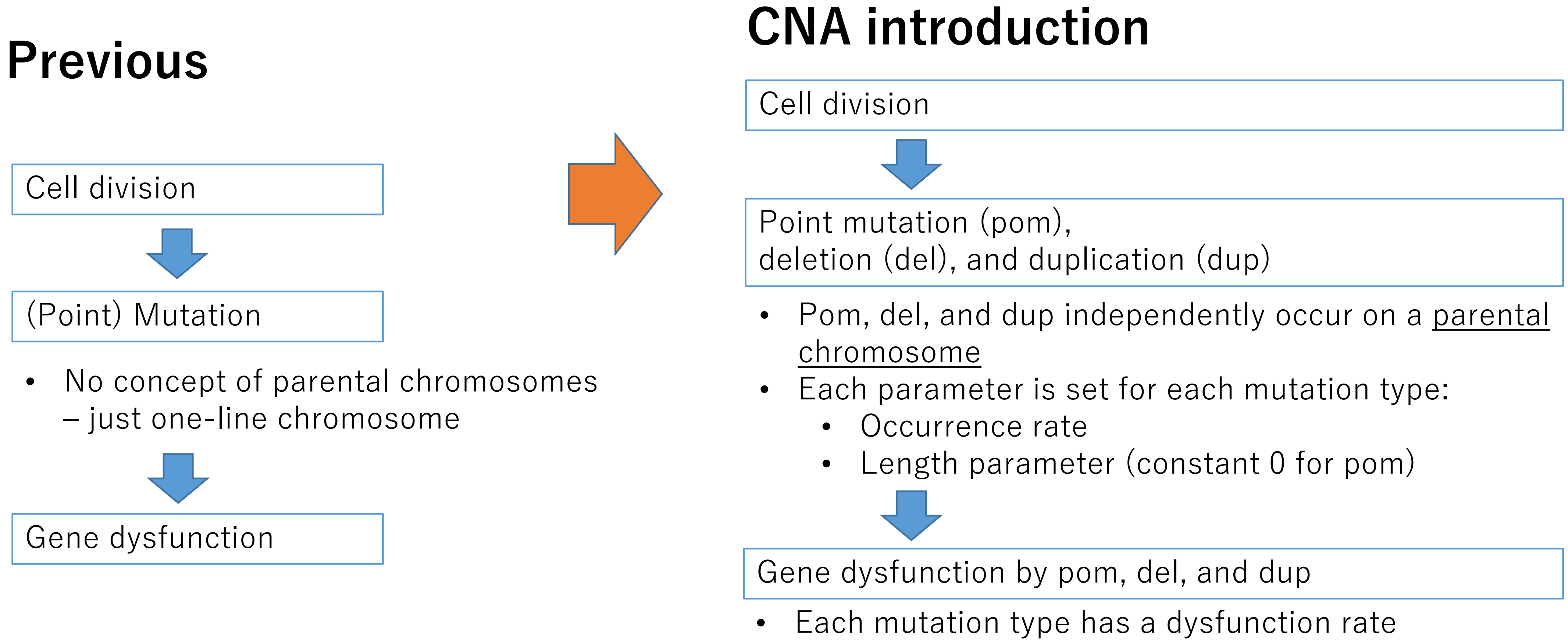

### Supplementary Figure 2 (contd.)

**(b)** Correction for pom

#### Previous

#### Current

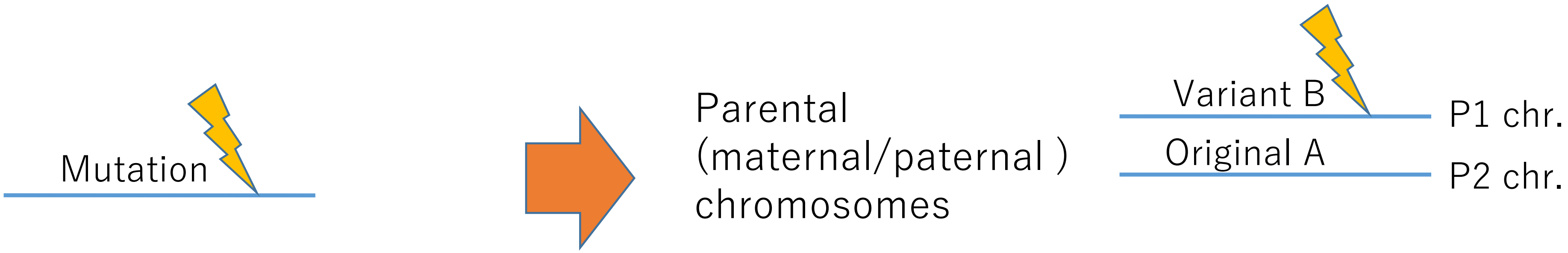


Previous:

1. No parental chromosome concept
   - A pom falls on a one-line chromosome
2. No conceptual distinction between exons and introns
3. Copy number of 2 and 100% tumor purity were assumed in VAF calculation

Current:

1. Introducing parental chromosomes
2. A pom is only in exons
3. CNA and the admixture rate of normal cells are introduced in VAF calculation

### Supplementary Figure 2 (contd.)

#### (c) Representative modes of CNA generation mechanisms

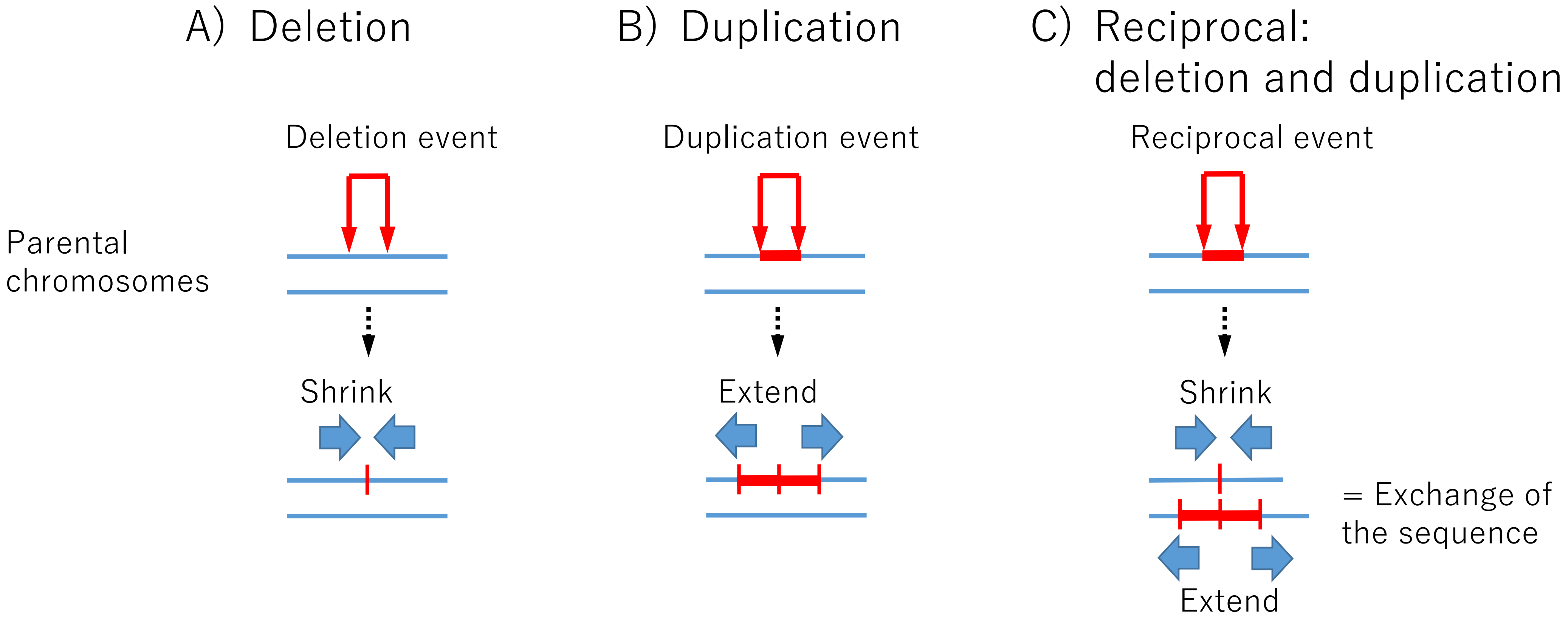

### Supplementary Figure 2 (contd.)

#### (d) Stochastic determination of the start and end points of a CNA

1. "**Fall**"
   - Stochastically determine the start position

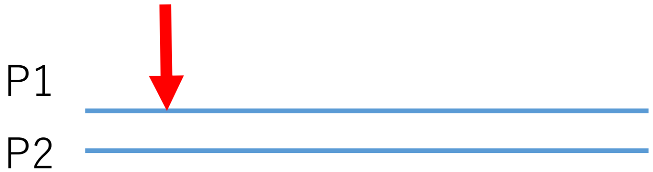


a. A start point occurs based on a **Poisson distribution** with the mean of the occurrence probability × target chromosomal length
   - The occurrence probability:
     For dup: *dup rate*
     For del: *del rate*

b. The position is determined based on a **uniform distribution**

2. "**Extend**"
   - Stochastically determine the length of a CNA
   - Then, the end position is automatically determined

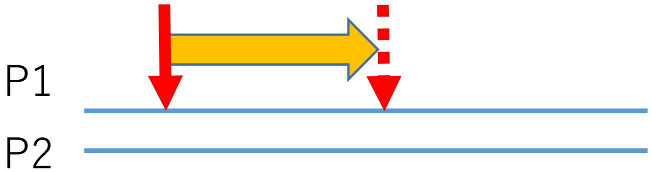


a. The length is determined based on an **exponential distribution**
   - For dup: *dup average length*
   - For del: *del average length*

### Supplementary Figure 2 (contd.)

**(e)** Gene dysfunction by CNA

- A point mutation may result in the dysfunction of a single gene.
- A CNA may result in the dysfunction of multiple genes.

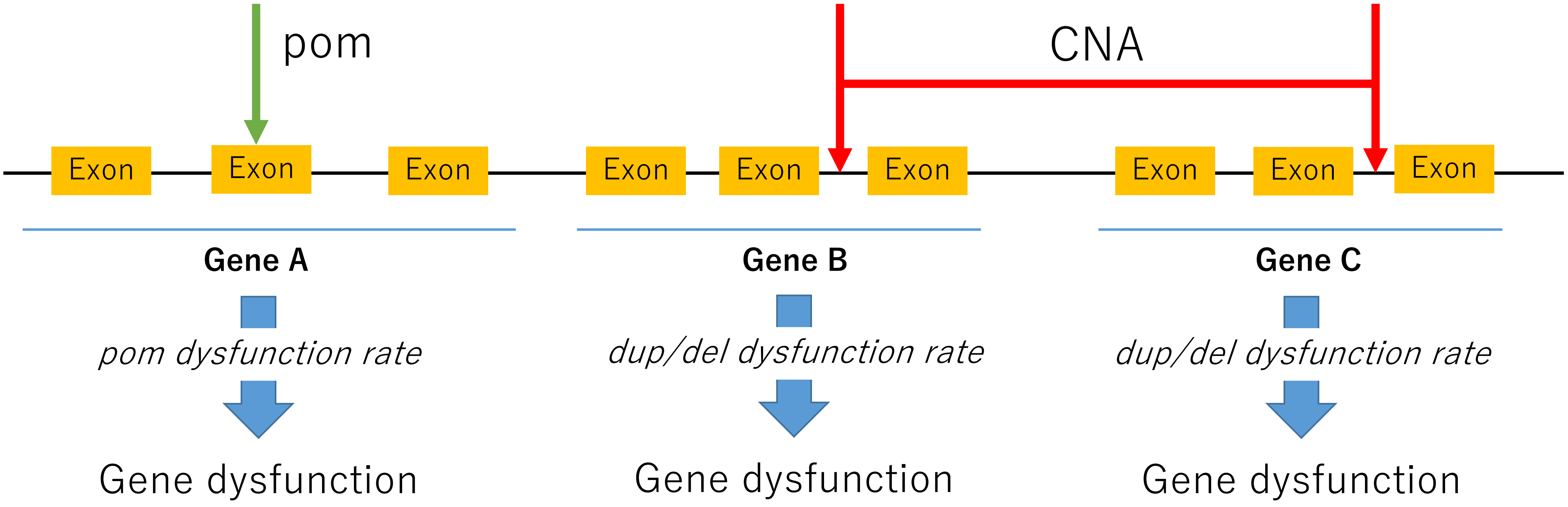

### Supplementary Figure 3

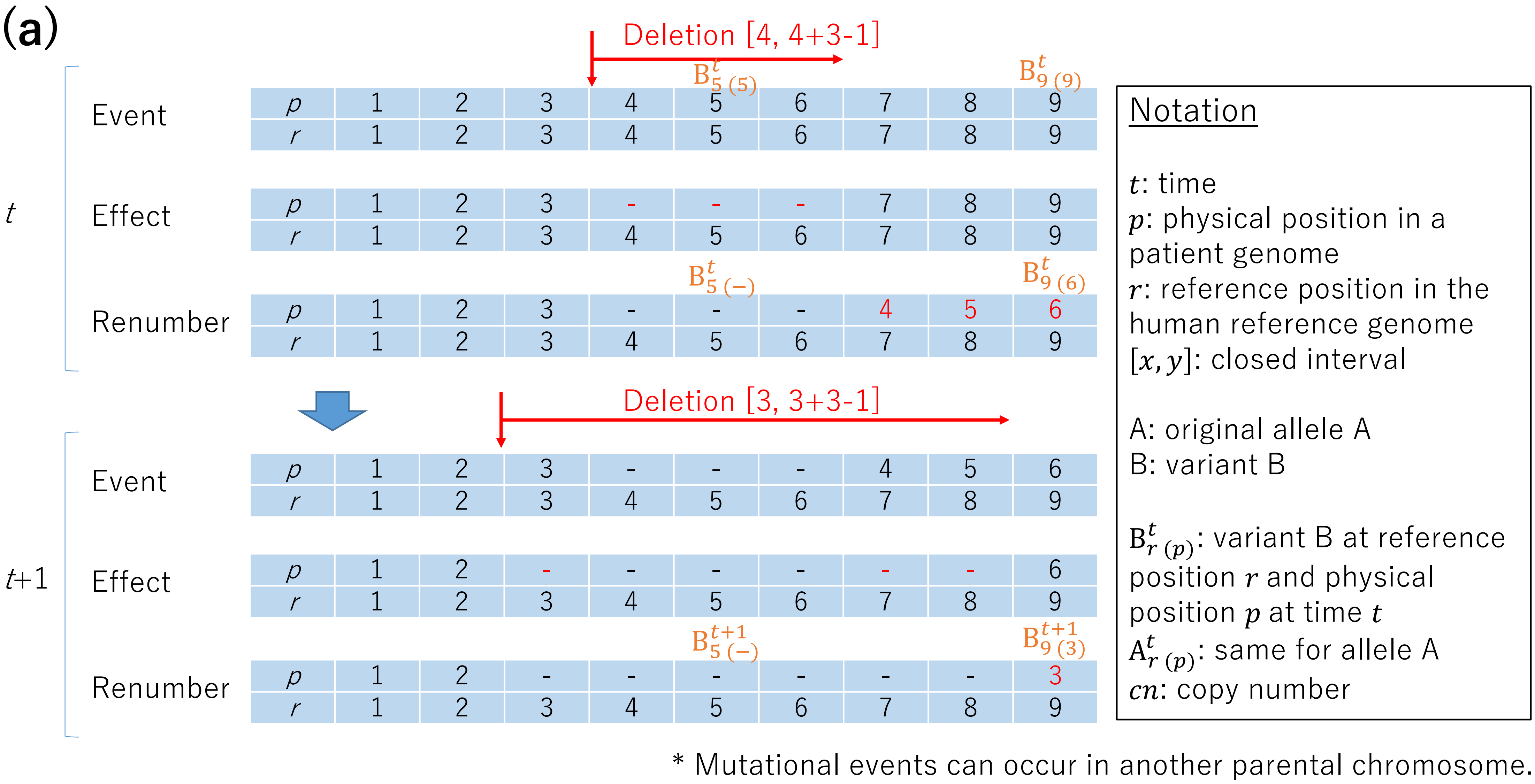

### Supplementary Figure 3 (contd.)

**(b)**

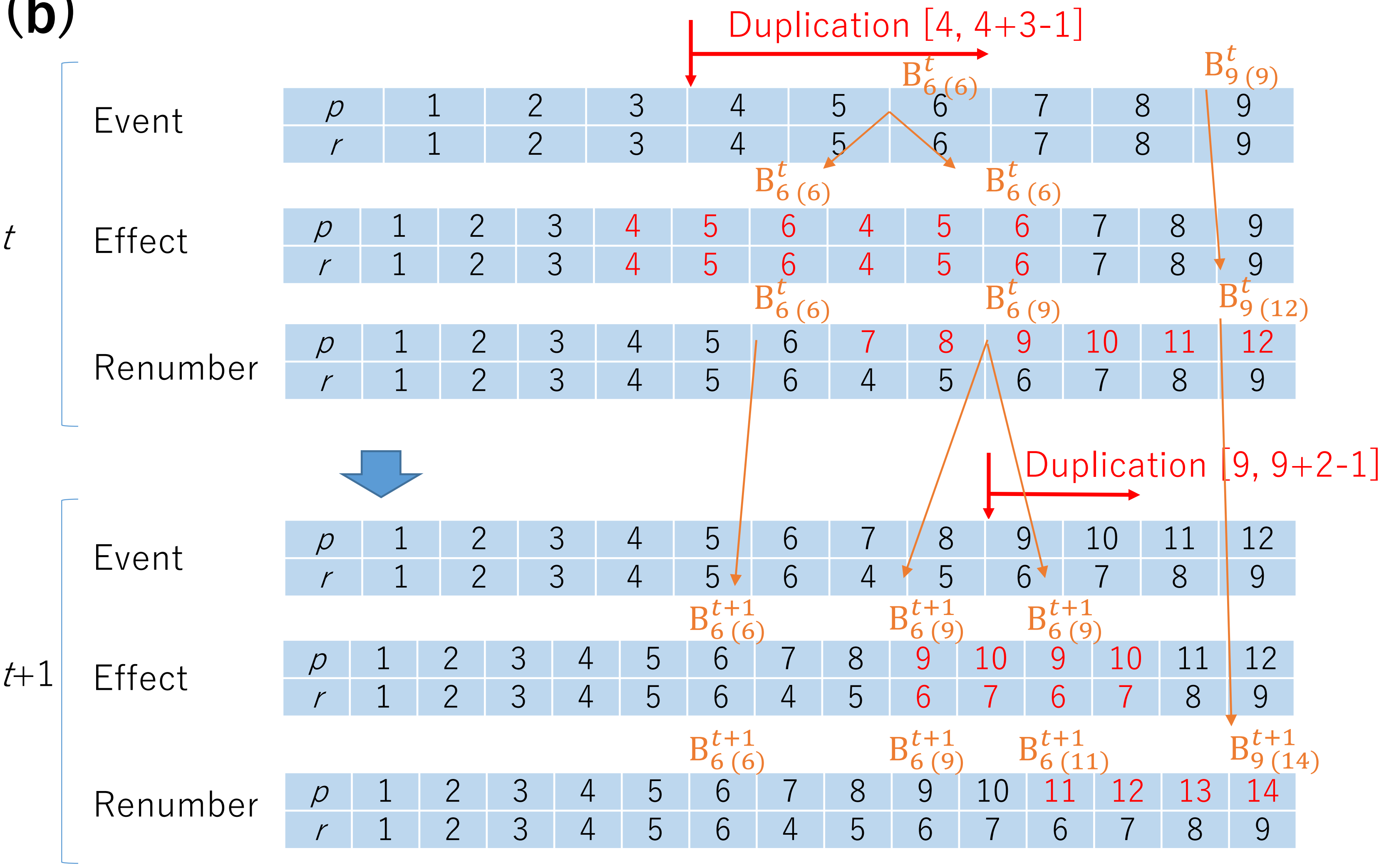

### Supplementary Figure 3 (contd.)

**(c)**

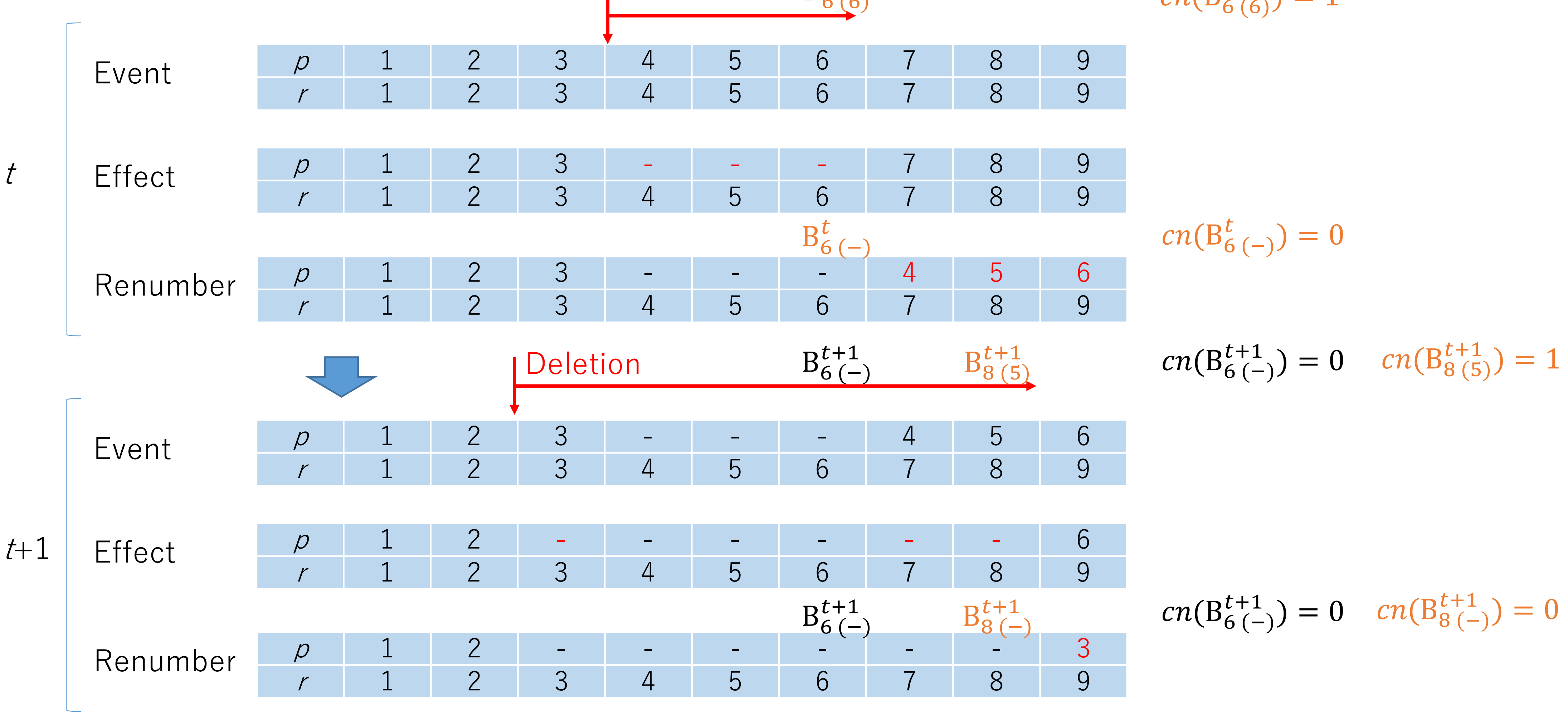

### Supplementary Figure 3 (contd.)

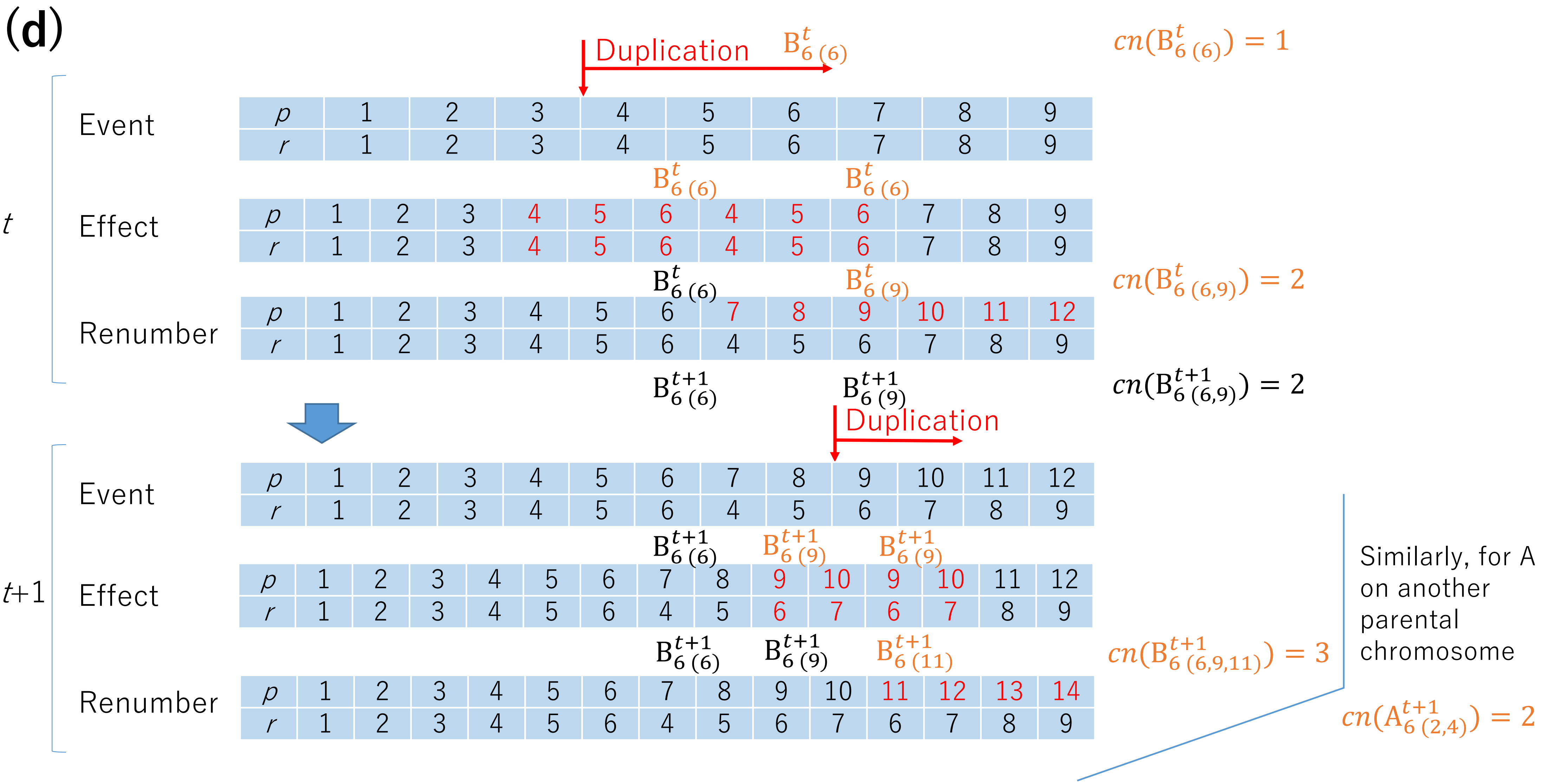

### Supplementary Figure 4

(a)

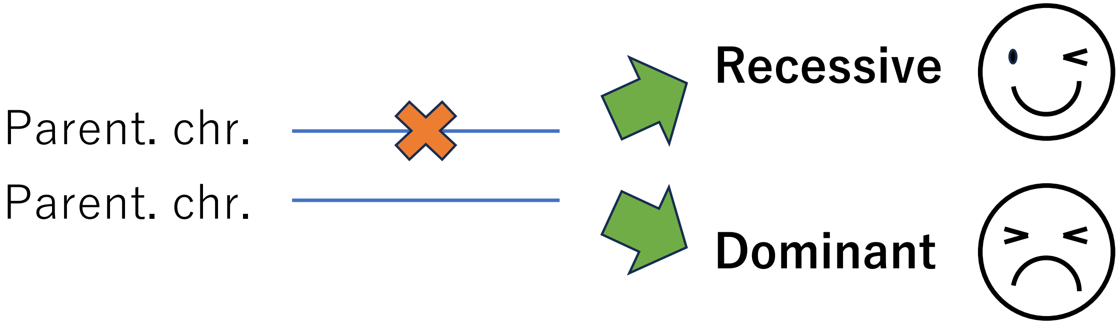


(b)

| | pom | del | dup |
|---|---|---|---|
| Oncogene | Dominant | - | Dominant |
| Suppressor | Recessive | Recessive | - |
| Dominant negative | Dominant | Recessive | - |

| | pom | del | dup |
|---|---|---|---|
| Oncogene | $u_{pom,o}$ | 0 | $u_{dup,o}$ |
| Suppressor | $u_{pom,s}$ | $u_{del,s}$ | 0 |
| Dominant negative | $u_{pom,s}$ | $u_{del,s}$ | 0 |

### Supplementary Figure 5

## (a)

$i$: position (site) index
$n$: copy number
$s$: subpopulation index
$\tau$: subpopulation fraction
$\rho$: proportion

A: original allele A
B: variant B
N: intact normal
SN: speckled normal
T: tumor

- $\rho_{\mathrm{NGS}} \sim \rho_{\mathrm{T}} + \rho_{\mathrm{SN}}$

where $\rho_{\mathrm{NGS}}$ is tumor purity measured from NGS

- $\rho_{\mathrm{pathology}} \sim \rho_{\mathrm{T}}$

where $\rho_{\mathrm{pathology}}$ is pathological tumor purity.

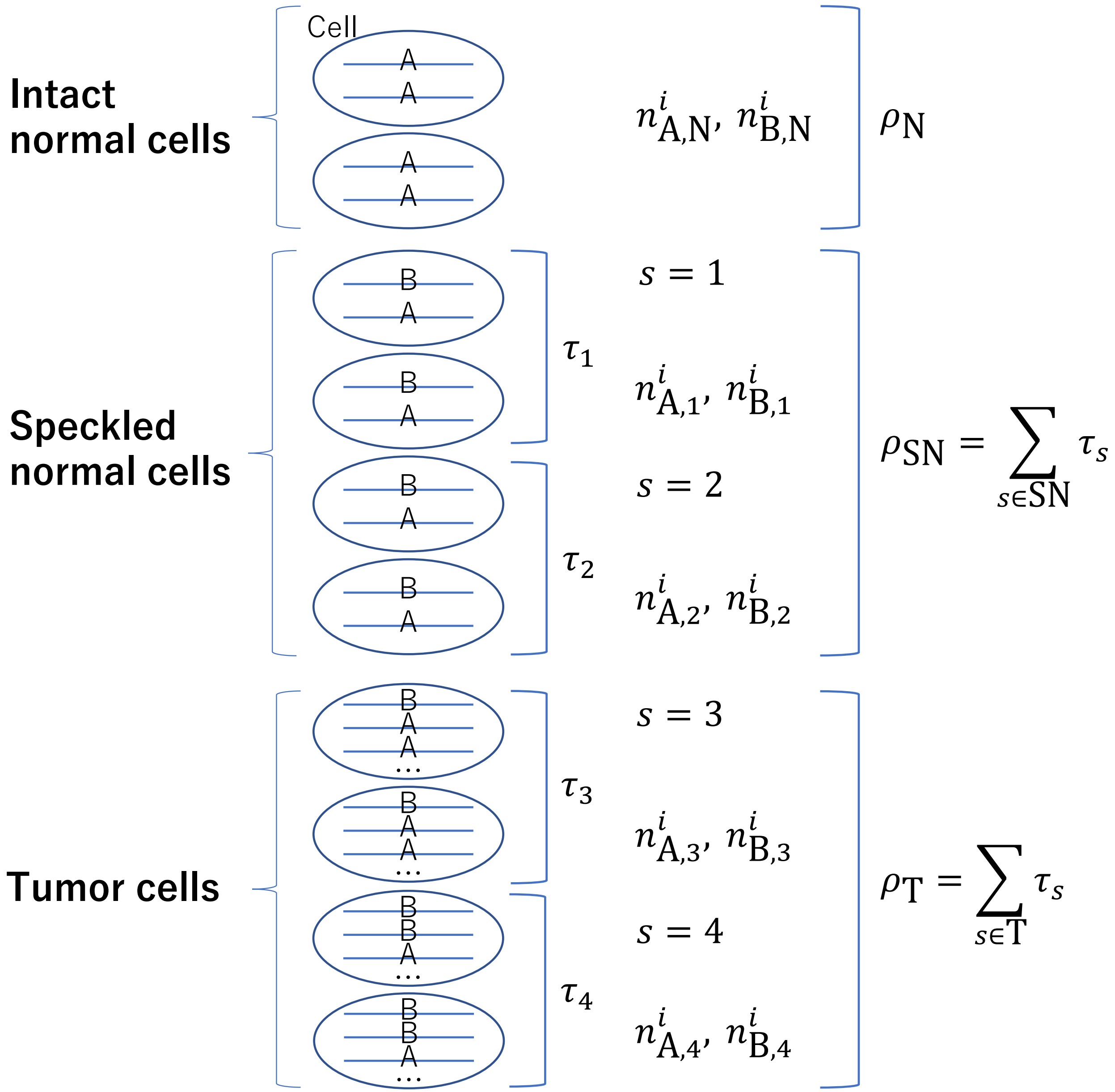

### Supplementary Figure 5 (contd.)

## (b)

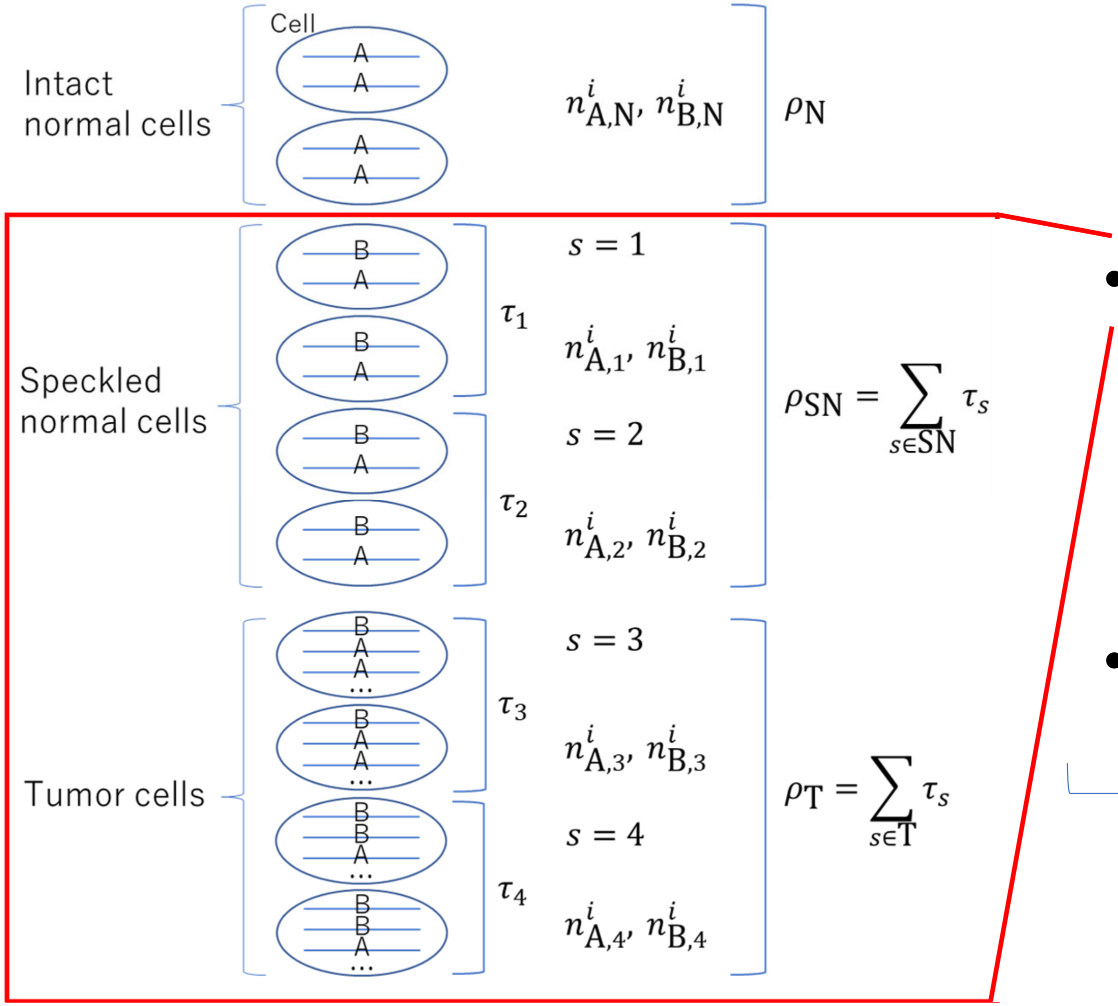


- $\rho_N$: parameter to be estimated
  - $\rho'_N$: arbitrarily given value, later checked by the fit between simulated and real VAFs
- $\lambda_s$: fraction in simulation
  - Fraction of a subpopulation comprising speckled normal and tumor cells in a simulation

$$\sum_{s=1}^{\#sp} \lambda_s = 1$$

- Then, scaled to a real fraction

$$\tau_s = (1 - \rho'_N)\lambda_s$$

VAF

### Supplementary Figure 6

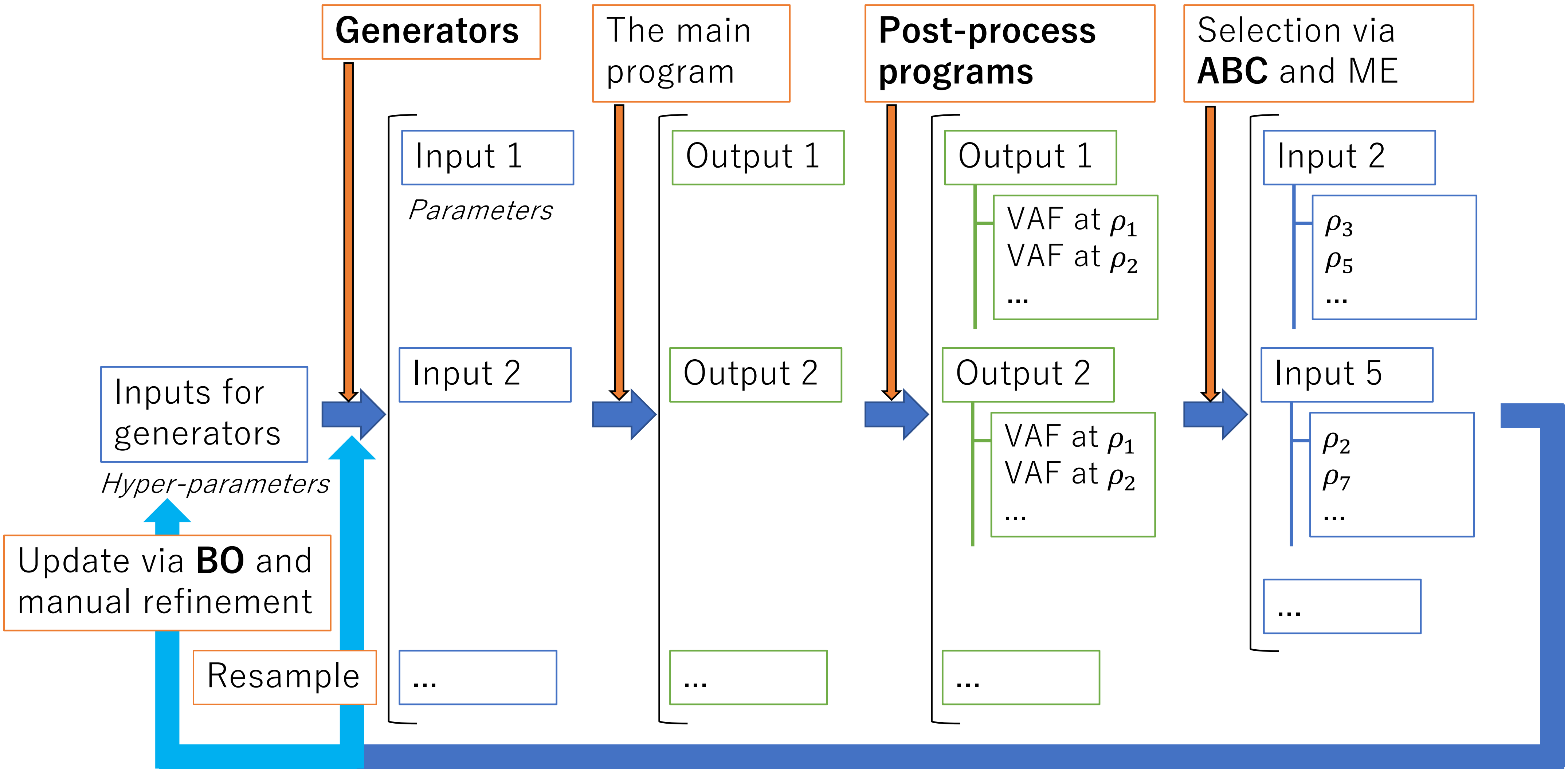

### Supplementary Figure 7

(a)

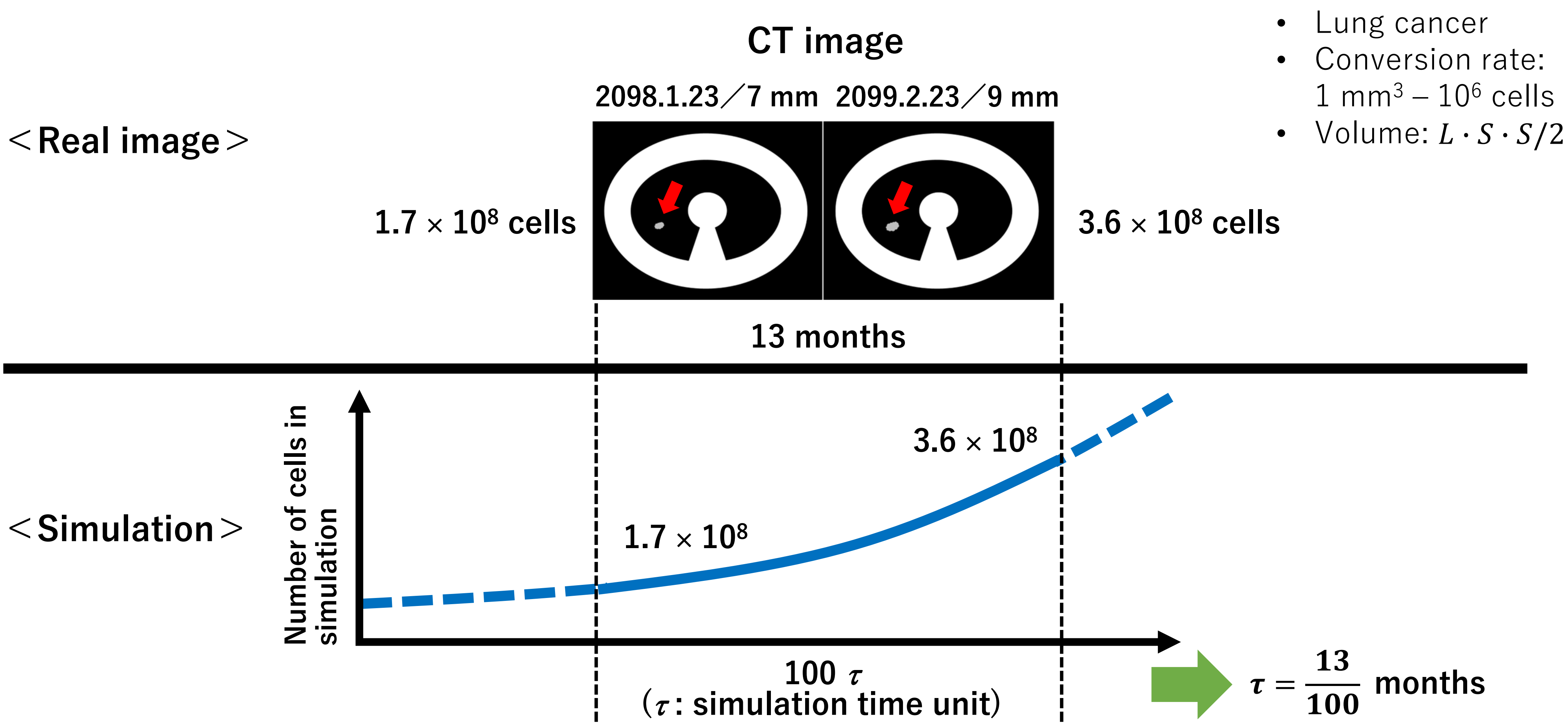

### Supplementary Figure 7 (contd.)

(b)

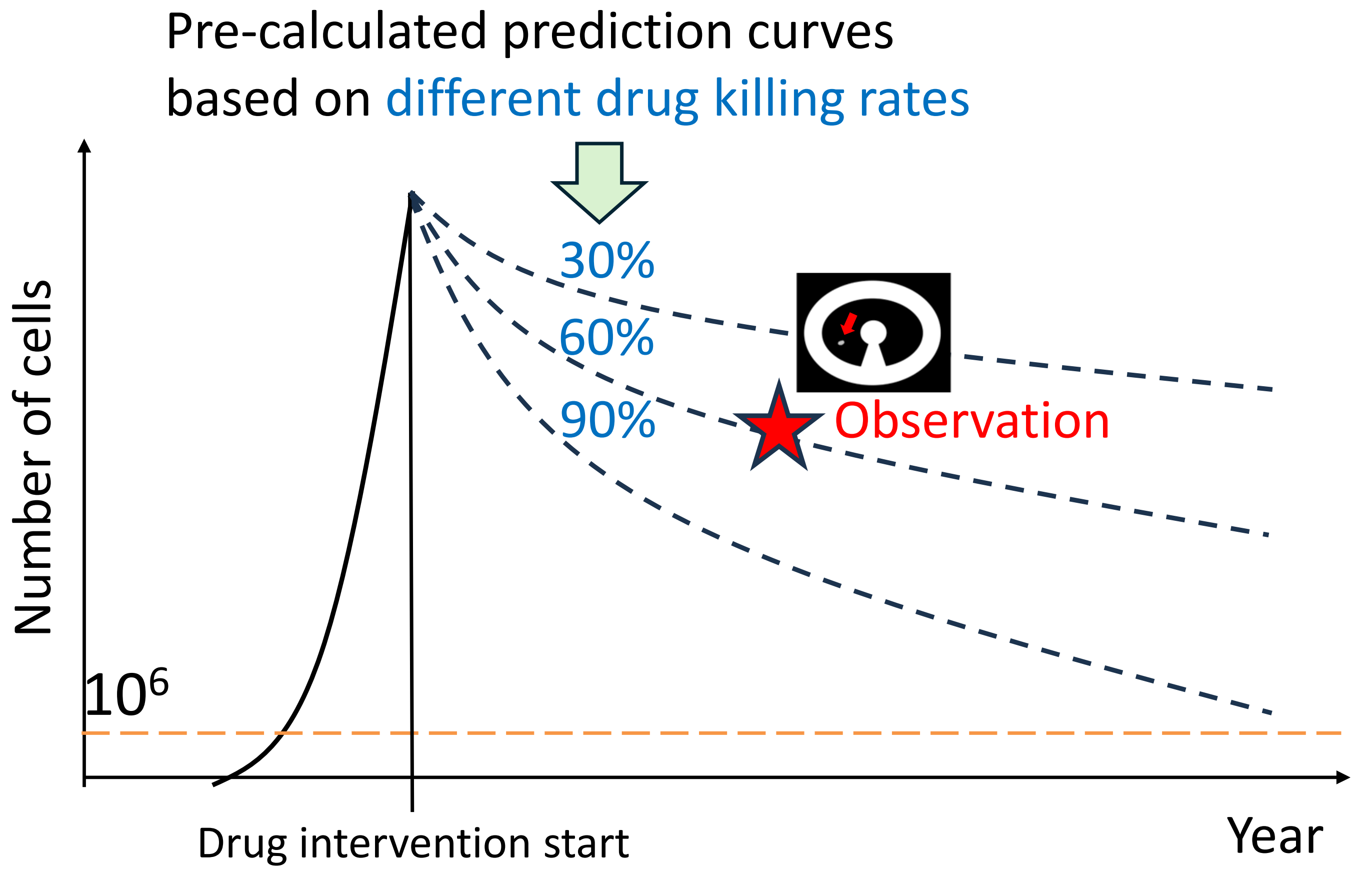

### Supplementary Figure 8

**(a)**

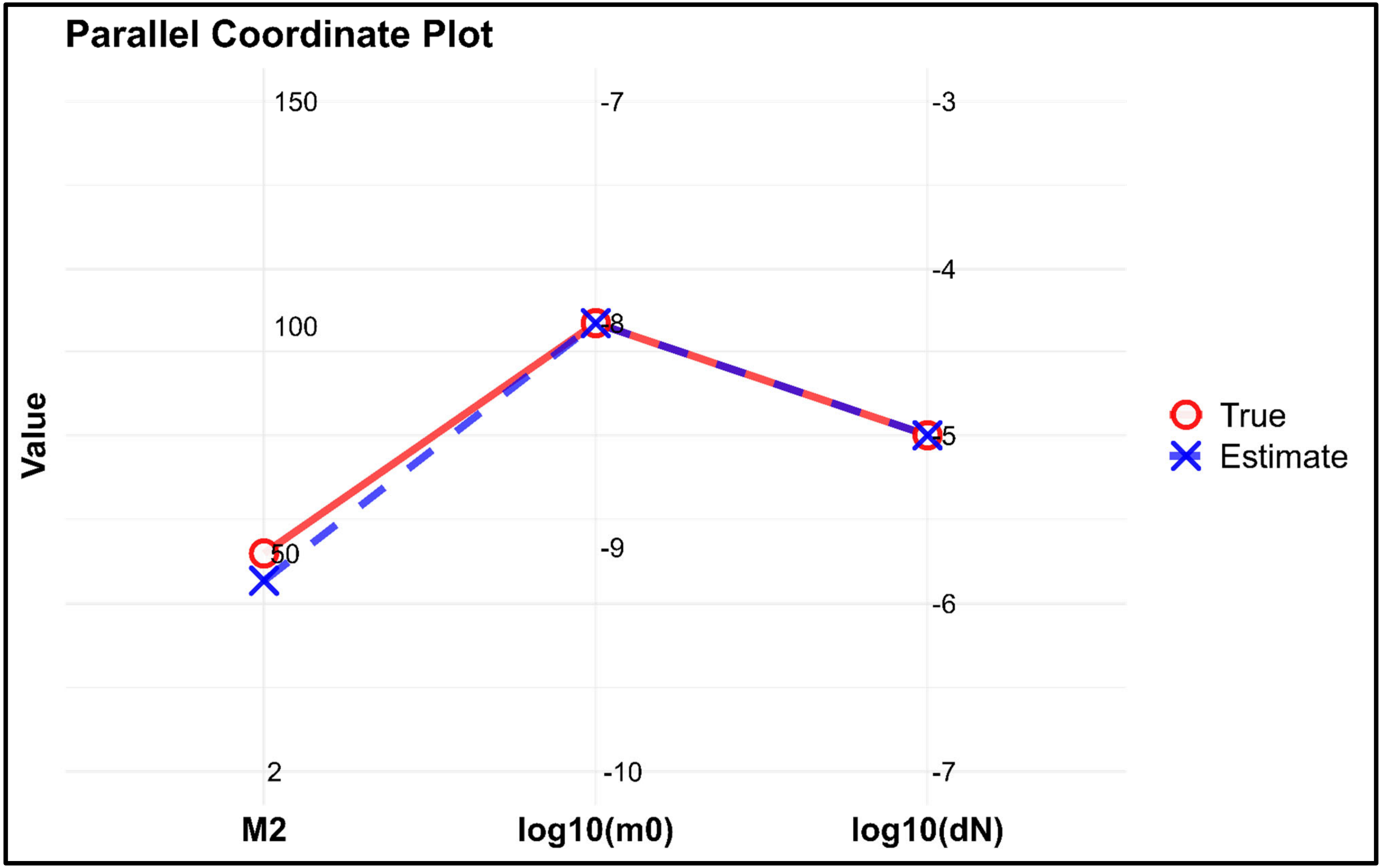

**Supplementary Figure 8 (cont'd)**

**(b)**

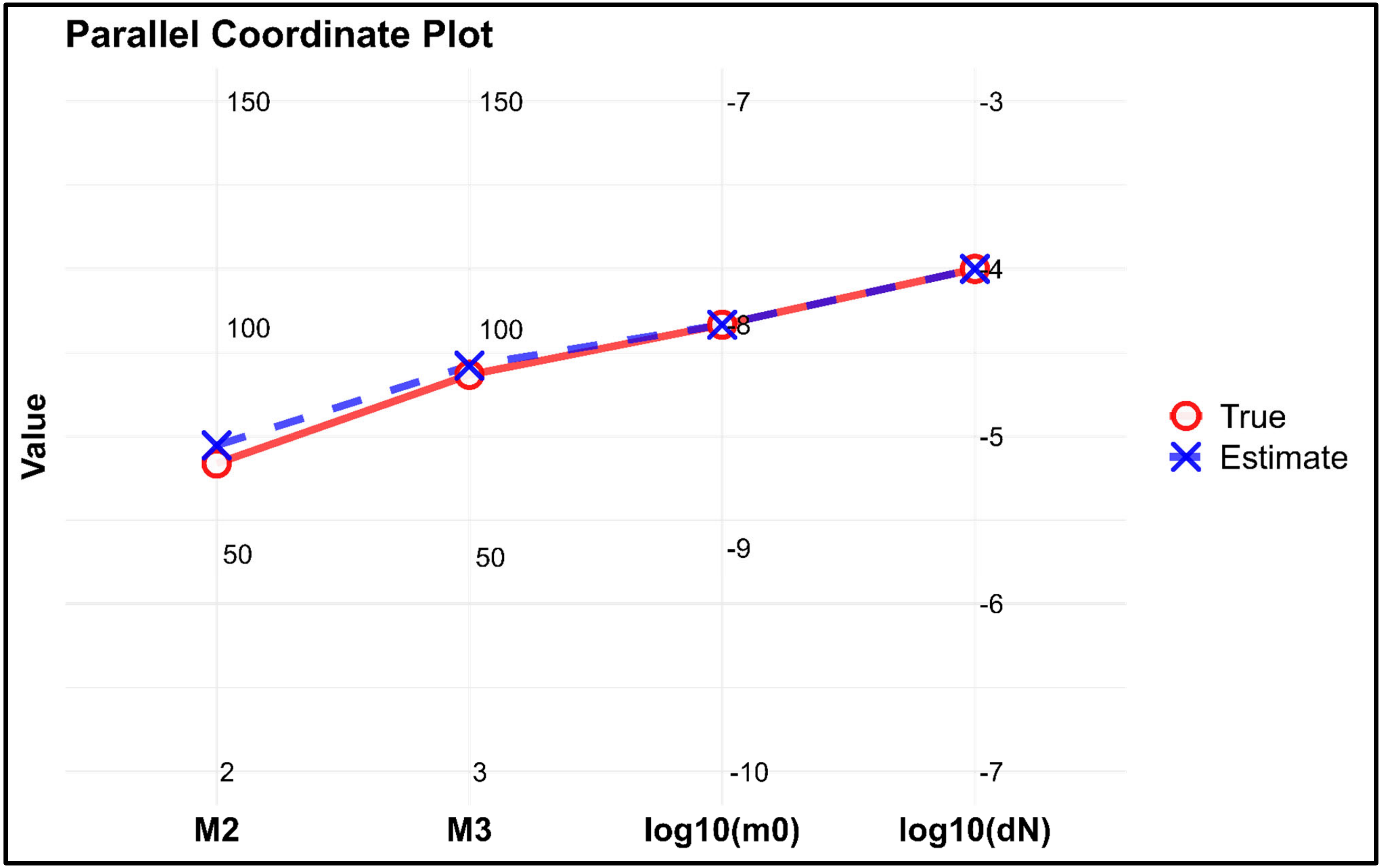

### Supplementary Figure 9

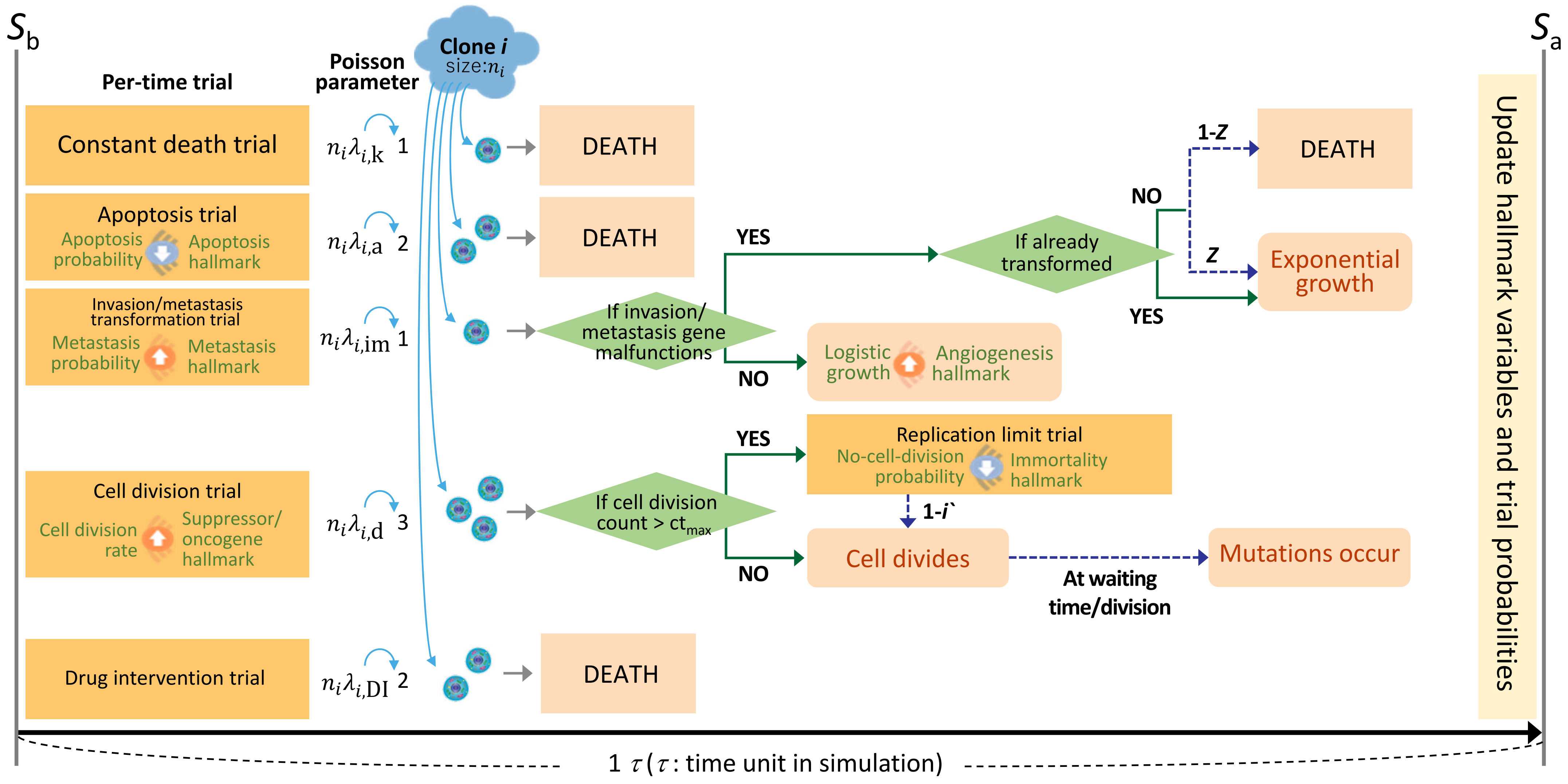
